# Microbial exposure across life reduces susceptibility of *Aedes aegypti* to Zika virus by enhancing blood digestion and limiting midgut cell infection

**DOI:** 10.1101/2022.11.10.516021

**Authors:** William Louie, Ana L. Ramírez, Rochelle Leung, Lindsey K. Mack, Erin Taylor Kelly, Geoffrey M. Attardo, Lark L. Coffey

**Affiliations:** Department of Pathology, Microbiology, and Immunology, School of Veterinary Medicine, University of California Davis, Davis, California, United States; Department of Entomology and Nematology, University of California, Davis, Davis, CA, United States

## Abstract

The worldwide expansion of mosquito-borne pathogens necessitates improved control measures, including approaches to reduce transmission by mosquito vectors. Reducing transmission is challenging because determinants of vector competence for viruses like Zika (ZIKV) are poorly understood. Our previous work established that *Aedes (Ae.) aegypti* larvae reared in environmental water containing microbes are less susceptible to ZIKV as adults compared to cohorts reared in laboratory tap water with fewer microbial species and lower microbial abundance. Here, we identify a process by which environment-derived microbes reduce susceptibility of *Ae. aegypti* for ZIKV. Provided that the midgut represents the first barrier to mosquito infection, we hypothesized that microbial exposure modulates midgut infection by ZIKV. Since mosquitoes live in water as larvae and pupae and then transition to air as adults, we also define the stage in the life of a mosquito when microbial exposure reduces ZIKV susceptibility. *Ae. aegypti* larvae were reared in water containing microbes and then treated with antibiotics during the pupal and adult stages, adult stage only, or provided no antibiotics at any stage. Vector competence was next evaluated in mosquitoes that ingested ZIKV-spiked bloodmeals. Antibiotic treated mosquitoes with reduced microbiota showed enhanced ZIKV infection rates in *Ae. aegypti* treated as both pupae and adults. Antibiotic treatment to disrupt microbes in pupal and adult mosquitoes also resulted in increased midgut epithelium permeability, higher numbers of ZIKV-infected midgut cells, and impaired bloodmeal digestion. Parallel control experiments with antibiotic-treated or gnotobiotic mosquitoes reared in laboratory water showed that the dysbiotic state created by antibiotic use does not influence ZIKV vector competence or midgut permeability and that more than the bacterial species in gnotobiotic mosquitoes is responsible for reducing ZIKV vector competence. *Ae. aegypti* with disrupted microbiota via antibiotic treatment as pupae and adults that ingested ZIKV in bloodmeals showed reduced expression of genes associated with bloodmeal digestion and metabolism relative to mosquitoes whose microbes were not reduced with antibiotics. Together, these data show that exposure to microbes throughout the life of *Ae. aegypti* restricts ZIKV infection by facilitating blood digestion and reducing midgut cell infection. Understanding the connections between mosquito microbiota, midgut physiology, and arbovirus susceptibility can lead to novel approaches to decrease mosquito transmission and will improve understanding of vector competence in environmental habitats containing microbes.

**Author Summary:** Mosquito-transmitted viruses like Zika continue to threaten human health. Absent vaccines or treatments, controlling mosquitoes or limiting their ability to transmit viruses represents a primary way to prevent mosquito-borne viral diseases. The role mosquito microbiota play in shaping transmission of Zika virus has been limited to association-based studies. Our prior work showed that *Aedes aegypti* mosquito larvae that develop in water containing bacteria are less susceptible to Zika virus compared to larvae reared in laboratory tap water with fewer numbers and species of bacteria. Here we identify a process that explains this association. Since mosquitoes are aquatic as larvae and pupae and terrestrial as adults, we also define the life stage when microbes need be present to reduce Zika virus susceptibility. We used antibiotics to reduce environmental water-derived microbes at pupal and adult or only adult stages and observed that microbial disruption via antibiotic treatment increases Zika virus infection and midgut permeability and impairs bloodmeal digestion. These findings advance understanding of microbiota-mosquito-virus interactions and further implicate microbes as a means to restrict virus infection of mosquitoes.

## Introduction

*Aedes (Ae.) aegypti* and *Ae. albopictus* mosquito-transmitted viruses, including dengue (DENV), chikungunya (CHIKV), and Zika (ZIKV), pose a massive burden on human health. Vector control represents the primary means to reduce the diseases they cause. Vector competence studies, which assess the capacity for mosquitoes to become infected with, develop disseminated infections, and transmit viruses in laboratory environments, are foundational to defining the species that maintain arbovirus transmission cycles. For *Ae. aegypti,* prior studies show that temperature (1,2) and geographic origin (3,4) are major determinants of ZIKV vector competence. Other factors, including the microbial environment of mosquito vectors, have been subject to much recent study (reviewed in Ferriera *et al.* (5)). The microbiota of *Aedes spp.* mosquitoes primarily derive from the environment (6–8), reside dominantly in the midgut (8,9), and can influence physiology and life history traits through nutritional supplementation (6,7,10,11) and immune stimulation (12–14). Prior studies from our group and others show that microbiota influence ZIKV competence for *Ae. aegypti* (15,16), and *Ae. albopictus* (17), and that these interactions are influenced by mosquito genotype (18). We discovered that when *Ae. aegypti* larvae reared in water with microbes sourced from cemeteries were exposed orally to ZIKV as adults, they became infected, developed disseminated infections, and transmitted virus less efficiently than cohorts where larvae were reared in laboratory tap water with fewer numbers and species of microbes (15). We also determined that the presence of microbes accelerates mosquito development. Another study demonstrated that disruption of microbiota via oral treatment of adult *Ae. aegypti* with antibiotics prior to exposure to ZIKV did not affect vector competence (19). Together, these findings suggest that the influence of microbiota on ZIKV vector competence may be specific to the mosquito developmental stage (larval, pupal, adult) during which the microbes are present. To understand whether reduced ZIKV vector competence after larval development in water containing microbes is conferred transstadially from larvae to pupae to adults, one goal of this study was to determine whether exposure to microbes throughout life is required to reduce ZIKV susceptibility in *Ae. aegypti*.

Microbes may modify vector competence of mosquitoes via multiple mechanisms, including to alter immune system activation, by direct antiviral or antiparasitic activity, or by affecting physiological processes, including blood digestion. Many of these processes occur in the mosquito midgut, which represents the first infection barrier for most vector-borne pathogens. Microbes are likely also involved in maintaining physical barriers that restrict access of an ingested virus or parasite to the mosquito midgut. The commensal bacterium *Serratia marcescens* increases susceptibility of *Ae. aegypti* for CHIKV by secreting a protein that digests membrane-bound mucins coating the gut epithelium, rendering the epithelium more accessible to the virus (20). The bacterium *Bacillus thuringiensis* svar. *israelensis* (Bti) expresses δ-endotoxins that create pores in midgut cells of larval mosquitoes, which is lethal and has long been a mainstay of larvicidal vector control (21,22). We hypothesized that disturbance in the microbiota of pupal and adult *Ae. aegypti* mosquitoes would decrease blood digestion, resulting in increased ZIKV susceptibility by giving the virus more time to access the midgut, since blood digestion is thought to occur concomitantly with virus infection of the midgut epithelium (23). We reared *Ae. aegypti* larvae in environmental water containing microbes. Half of the experimental cohorts were administered antibiotics in water during the pupal stage to reduce microbes. Some cohorts of adults were also administered antibiotics in sugar water prior to exposure to ZIKV-spiked bloodmeals. Since antibiotics create a dysbiotic environment, may not eliminate resident antibiotic-resistant microbiota (24), and can alter cellular metabolism and transcription as described in mammalian cells (25–28), *Drosophila* (29), and *Anopheles* (30) in ways that could also impact ZIKV infection in *Ae. aegypti*, we first verified that antibiotic treatment did not modify midgut integrity or ZIKV vector competence. We also generated gnotobiotic mosquitoes by colonizing larvae with an *Elizabethkingia spp.* strain; this bacterial species was not sufficient to modulate vector competence, suggesting that many species are required. Next, we measured physiological differences in the midgut in blood engorged mosquitoes across cohorts treated with or without antibiotics at different life stages. Microscopy was used to quantify ZIKV-infected midgut cells. The magnitude and rate of bloodmeal digestion was measured by quantifying hemoglobin levels over time. Since the ways microbes modify vector competence are not fully understood, RNA sequencing (RNA-seq) was used to identify differential gene expression in cohorts of *Ae. aegypti* treated with or without antibiotics at different life stages and in response to blood with or without ZIKV. We observed that *Ae. aegypti* given antibiotics as pupae and adults are more susceptible to ZIKV infection compared to those not given antibiotics. Microbe reduction by antibiotic treatment of pupae and adults also increased midgut permeability, the number of ZIKV-infected midgut cells, and impaired bloodmeal digestion. Antibiotic treatment itself did not reduce ZIKV vector competence or affect midgut permeability, ruling out toxic or physiologic effects from the antibiotic cocktail used. Gene ontology analyses of ZIKV-exposed mosquitoes revealed enrichment of PM-associated genes, with more genes related to blood protein metabolism upregulated in mosquitoes harboring intact microbiota. Together our results show that exposure to microbes throughout the life of *Ae. aegypti* facilitates blood digestion and reduces ZIKV midgut cell infection.

## Materials and Methods

### Biosafety

All ZIKV infection experiments were conducted in a biosafety level 3 (BSL3) laboratory and were approved by the University of California, Davis (UC Davis) Institutional Biosafety Committee under Biological Use Authorization #R1863.

### Mosquito sources

*Ae. aegypti* mosquitoes colonized from field collections from 2 locations in California were used. Mosquitoes were field-collected in 2016 as larvae (Los Angeles) and eggs (Clovis). Mosquitoes from both locations were maintained for generations numbers noted in Figures representing F_20_, F_22,_ or F_26_, at 28°C, 80% humidity, and a 12 h:12 h light:dark (L:D) cycle with 10% sucrose provided to adults and periodic presentation of sheep blood to stimulate egg laying, as described in our previous study (15).

### Mosquito generation and antibiotic treatments

*Microbe source and preparation*. Microbes from water collected outdoors in cemeteries were added to water in which *Ae. aegypti* larvae were reared, using an approach as in our previous study (15). Briefly, 1 liter (L) of collected water was filtered through a 1 mm mesh to remove large debris, centrifuged at 3000 g to pellet microbes, washed 3 times with sterile 1X phosphate buffered saline (PBS, Thermo Fisher Scientific, Emeryville, CA), and frozen in glycerol at-80°C. Microbe aliquots were thawed at room temperature and placed in laboratory tap water prior to addition of mosquito eggs.

*Larval rearing.* Mosquito eggs were hatched in 1 L deionized (diH_2_O) water spiked with 1 mL of microbes from a glycerol stock aliquot. Approximately 1500 larvae were reared in three pans at a density of ∼500 larvae/pan. Each pan contained an agarose plug that was prepared by mixing 1% agarose (Sigma-Aldrich, St. Louis, MO) with pulverized fish food (final concentration of 100 g/L, or 10% [Tetra, Melle, Germany]) and rodent chow (final concentration of 80 g/L, or 8% [Teklad Global 18% Protein Rodent Diet, Envigo, Indianapolis, IN]), which was autoclave-sterilized before casting into 12-well plates. One to 2 plugs were added to larval pans every other day until day 15 post-hatch. Larval trays were kept in environmental growth chambers (Binder, Bohemia, NY) at 28°C, 80% humidity, and a 12 h:12 h L:D cycle.

*Antibiotic treatments to pupae and adults*. Pupae or adults were divided into three treatments:1) mosquitoes exposed to antibiotics (Abx) both as **p**upae and **a**dults (AbxPA), 2) mosquitoes exposed to antibiotics as **a**dults (AbxA) only, and 3) mosquitoes not exposed to antibiotics (No Abx), where the latter represents the control group. First, pupae were picked and randomly divided into 2 batches each day. The AbxPA batch of pupae was washed in diH_2_O containing an antibiotic cocktail consisting of 50 U/mL each of penicillin, streptomycin, and kanamycin (Sigma-Aldrich, St. Louis, MO) for 10 minutes before transfer to another water vessel containing fresh antibiotic cocktail. This concentration is in the middle of the range of doses used previously in similar studies with *Ae. aegypti* (20) and *An. stephensi* (31). The other batch of pupae, from which the AbxA and No Abx groups were derived, was transferred to a separate cage without washing or antibiotics. Adults in the AbxPA group were maintained on the antibiotic cocktail dissolved in 10% sucrose (Thermo Fisher Scientific, Emeryville, CA) *ad libitum*. Adults in the other batch were initially maintained on filter-sterilized 10% sucrose without antibiotics *ad libitum* until eclosion was complete. Once adults reached a density of ∼500, females were aspirated out and transferred in cohorts of 80-120 mosquitoes to 32-ounce plastic containers (Amazon, Seattle, WA) with mesh lids. At this point, the second non-antibiotic treated batch was further divided into two treatments:AbxA was provided 10% sucrose *ad libitum* with the antibiotic cocktail, and No Abx was fed 10% sucrose *ad libitum* without the antibiotic cocktail. As a control for toxicity or physiological effects stemming from antibiotic treatment, we also reared mosquitoes in laboratory tap water (conventional) with or without antibiotics, which were provided as described above, to pupal and adult mosquitoes (conventional + Abx). Adult mosquitoes were maintained at 28°C, 80% humidity, and 12 hour:12 hour L:D cycle throughout the experiments. All mosquitoes were housed in the same incubator. Antibiotic treatment experiments with conventional mosquitoes and experiments with mosquitoes provided environmental microbes each performed once.

### Gnotobiotic mosquito generation

An *Elizabethkingia spp.* strain isolated from colonized *Ae. aegypti* from Los Angeles was cultured in LB broth at 37°C for 24 hours prior to inoculation into larval flasks. *Ae. aegypti* eggs attached to their egg papers cut into strips were surface sterilized by submersion in 1% bleach for 10 minutes. After 10 minutes, bleach was removed by vacuum filtration of eggs through 1000 mL of diH_2_O and a 0.22 μm filter (Genesee Scientific, San Diego, CA). Eggs were then washed twice in 70% ethanol and allowed to dry. To prepare larval flasks, 50 mL of diH_2_O and one larval nutrient plug were added to four T75 tissue culture flasks. Paper strips containing ∼200 eggs were placed into each tissue culture flask and spiked with 1 mL of 1.4 x 10^8^ CFU/mL *Elizabethkingia* culture before sealing the lid and incubating at 28°C with 80% humidity and 12 h:12 h L:D cycle for 5 days. When pupae developed, lids of the tissue culture flasks were removed, and a sterilized fabric sleeve feeding into a makeshift plastic bucket with a mesh lid was taped around the opening of the flask to allow adults to emerge into the new holding container. Individual mosquitoes of each life stage were sampled, homogenized, and cultured on LB plates to confirm *Elizabethkingia* colonization and the absence of other contaminating bacteria, which was verified by colony morphology.

### Bacterial quantification by 16S qPCR

We measured the efficacy of antibiotic treatment for removing bacteria from AbxA and AbxPA groups and to verify dominance of *Elizabethkingia* in gnotobiotic mosquitoes by performing 16S qPCR. AbxA and AbxPA larvae, pupae, and adults 3-5 days post-eclosion (dpe) were evaluated. L4 larvae and pupae before antibiotic treatment were also included. Whole adult mosquitoes in the gnotobiotic group were evaluated and conventional and conventional+ Abx mosquitoes were included as comparators. DNA from 5 L4 larvae, 5 pre-treatment pupae, 5 No Abx, 10 AbxA, 10 AbxPA, 8 gnotobiotic, 10 conventional, and 10 conventional + Abx individuals was extracted with the Quick-DNA Tissue/Insect Microprep Kit (Zymo Research, Irvine, CA, USA), according to the manufacturer instructions and eluted in 30-40 µL elution buffer. Extracted DNA was PCR-amplified in the V3-V4 (32) hypervariable region of the 16S rRNA gene, as previously described (15,32). FASTQ files were processed using a standard DADA2 pipeline(33) (package version 1.22.0) into phyloseq (package version 1.38.0)(34). Sequences were assigned to ASVs, and samples were filtered by prevalence (ASVs at > 0.1% abundance), known taxonomy (filtered unknown taxa), and presence in the negative control (ASVs also present in the negative control were excluded).Quantification of bacteria was performed by SYBR Green Real-Time PCR (Thermo Fisher Scientific, Emeryville, CA) to amplify the 16S rRNA gene. No Abx, AbxA, and AbxPA samples were assayed in technical duplicates; gnotobiotic, conventional, and conventional + Abx were assayed once. 16S quantities were normalized to an *Ae. aegypti* reference ribosomal protein S17 gene (RPS17) and reported as a 16S:RPS17 ratio (35). The rationale for reporting the ratio instead of absolute values was to enable comparison of relative measurements across mosquito treatment groups to verify that antibiotic treatment reduces bacteria in mosquitoes.

### Hemolytic activity

Hematophagous bacteria can directly lyse blood cells and confound hemoglobin digestion measurements. Bacterial isolates from adult female *Ae. aegypti* were screened for hemolytic activity via culturing on agar plates. Individual mosquitoes were homogenized (TissueLyser, Retsch, Haan, Germany) in 500 µL PBS and 40 µL of supernatant was plated on either R2A agar (Sigma-Aldrich, St. Louis, MO) or sheep blood agar plates (Biological Media Services, UC Davis) for 3 days at 37°C. Up to 10 colonies from each mosquito were streaked on media plates 2-3 additional times to generate axenic cultures. Resulting colonies were streaked on sheep blood agar at both 30°C and 37°C and screened for clearing zones around colonies over the course of 3 days. The viability of this assay was tested in-house with hemolytic *Streptococcus spp.* isolates from the UC Davis School of Veterinary Medicine.

### Virus source and titrations

We used ZIKV strain BR15 (Brazil 2015, SPH2015, GenBank Accession # KU321639), the same strain we used previously for mosquito infection experiments (15). This virus strain was originally isolated from human serum from a patient in Brazil in 2015 and passaged 3 times in Vero cells (CCL-81, ATCC, Manassas, VA) before freezing in stocks. Stocks were titrated on Vero cells prior to bloodmeal presentation to approximate stock titers, and ZIKV-spiked sheep blood was also titrated after bloodfeeding to confirm the dose presented in each bloodmeal. For titrations, serial 10-fold dilutions of cell culture supernatant containing virus stock or ZIKV-spiked blood were inoculated into multi-well plates containing confluent Vero cells at one dilution per well and incubated for 1 hour at 37°C in 5% CO_2_ with rocking every 10 minutes. After incubation, each well was filled with an agar overlay (0.5% agarose [Thermo Fisher Scientific, Emeryville, CA] in Dulbecco’s modified Eagle’s medium [DMEM, Thermo Fisher Scientific, Emeryville, CA], + 2% fetal bovine serum [FBS, Genesee Scientific, San Diego, CA] + penicillin/streptomycin [Thermo Fisher Scientific, Emeryville, CA]). The cells were incubated for 1 week at 37°C in 5% CO_2,_ then fixed with 4% formalin (Thermo Fisher Scientific, Emeryville, CA). Fixed cells were stained with 0.025% crystal violet (Thermo Fisher Scientific, Emeryville, CA) in 20% ethanol and visualized on a light table to count plaques. ZIKV bloodmeal titers are represented as plaque-forming units (PFU) per mL of blood.

### ZIKV vector competence experiments

Stock ZIKV in DMEM, or DMEM with no virus as a control, were diluted in 10-fold increments with fresh heparinized sheep blood (HemoStat Laboratories, Dixon, CA) to achieve ZIKV titers of 10^6^ PFU/mL. Bloodmeals were presented to female *Ae. aegypti* at 3-5 days post-eclosion in cohorts of 80-120 per container, at 2-3 containers per treatment. Mosquitoes were sucrose-starved 18-24 hours before the presentation of bloodmeals. Mosquitoes were offered bloodmeals for 60 minutes through a collagen membrane rubbed with an artificial human scent (BG-Sweetscent mosquito attractant, Biogents USA) and heated to 37°C in a membrane feeder (Hemotek Limited, Blackburn, United Kingdom). Bloodmeals were presented to mosquitoes inside a plexiglass glove box containing an open bottle of boiling water to maximize humidity. After feeding, mosquitoes were immobilized with CO_2_ for 5 seconds and then immobilized on a chill table (Bioquip (R.I.P.), Compton, CA, USA) to assess their bloodfeeding status using a microscope at 10X magnification. Fully engorged females (18-45 per treatment for each experiment) were sorted into fresh plastic containers at a density of 10-20 mosquitoes per container and held at 28°C with 80% humidity and 12 h:12 h L:D cycle for 14 days. No Abx cohorts were provided 10% sucrose *ad libitum,* while AbxPA, AbxA, and conventional + Abx cohorts were maintained on 10% sucrose with the antibiotic cocktail *ad libitum*.

At 14 days post-bloodfeed (dpf), surviving mosquitoes were CO_2_-anesthetized and immobilized on a chill table. Bodies, legs/wings, and saliva were collected to assess infection, dissemination, and transmission rates, respectively. Legs/wings were removed before collection of expectorate into capillary tubes containing PBS by forced salivation, using an established approach (36). To minimize cross-contamination, surgical tools were washed once in CaviCide® (Metrex, Orange, CA) and twice in 70% ethanol between dissections. Each capillary tube was placed in a 1.5-mL tube containing 250 µL PBS and pulse-centrifuged at 8,000 g to recover saliva. Bodies and legs/wings were placed into 2-mL tubes containing 500 µL PBS and a 5-mm glass bead (Thermo Fisher Scientific, Emeryville, CA), then homogenized at 30 hertz for 5 minutes in a TissueLyser (Retsch, Haan, Germany). Viral RNA was extracted using a MagMax Viral RNA Extraction Kit (Thermo Fisher Scientific, Emeryville, CA), into 60 µL elution buffer following the manufacturer’s recommendations.

Detection and quantification of viral RNA in mosquito tissues and saliva were performed by quantitative reverse transcription polymerase chain reaction (RT-qPCR). The rationale for using RT-qPCR was to enable rapid and high throughput analyses, because the limit of detection (LOD) is lower than for titrations of volume limited samples, and to reproducibly enable comparison of relative measurements across mosquito treatment groups. Furthermore, our prior studies with ZIKV strain BR15 in Vero cells showed high reproducibility and a ZIKV RNA:PFU ratio of 100-2000 (37). We used a TaqMan Fast Virus 1-Step Master Mix and a ZIKV-specific primer set (ZIKV 1086F/1162c, probe:ZIKV 1107-FAM) using established methodologies (36,37). Cycle threshold (Ct) values from the RT-qPCR were converted to RNA genome copies using standard curves established with ZIKV RNA of known concentration. Each sample was assayed in technical duplicates and converted to copies per mosquito before calculating the average value. The LOD was calculated from the standard curve linear regression line where the Ct value = 40. Samples without a detectable Ct of less than 40 were reported at the LOD. Infection rates are reported as the number of ZIKV-positive individuals divided by the number that ingested ZIKV-spiked blood. Dissemination rates are reported as the number of ZIKV-positive legs and wings divided by the number of ZIKV-infected individuals. Transmission rates are reported as the number of ZIKV-positive saliva samples divided by the number individuals with disseminated ZIKV in legs and wings.

### Midgut permeability assessments

A low molecular weight dextran-FITC (MW 4000, Sigma-Aldrich, St. Louis, MO) was co-spiked into ZIKV-bloodmeals and used to measure midgut permeability in No Abx, AbxA, AbxPA, conventional, conventional + Abx, and gnotobiotic groups that were evaluated within 1 hour after bloodfeeding. Permeability was assessed by measuring the fluorescent intensity of dextran-FITC inside versus outside at three locations in the midgut for each mosquito for a total of 5 mosquitoes per treatment. A stock solution of 20 mg/mL dextran-FITC dissolved in PBS was spiked into sheep blood at a 1:20 dilution to achieve 1 mg/mL, the concentration used in a previous study (38). Dextran-spiked blood mixed with 10^5^ PFU/ml ZIKV and presented to mosquitoes as described above for vector competence assessments. Blood-engorged mosquitoes were visualized by fluorescence microscopy to detect FITC in tissues. Samples were covered in foil for all post-dissection steps to minimize photobleaching. Quantification of FITC fluorescence was then performed on Fiji (ImageJ) (39) by measuring intensity relative to background by subtracting fluorescence of an equal-sized region outside the sample area. Fluorescence intensity ratios between the gut lumen and hemocoel were calculated at three randomly drawn areas of equal size around the gut perimeter, then averaged together. Each average was also conducted across three slices per sample to minimize focal plane variation in fluorescence. At least 4 mosquitoes per group were included in each permeability assessment.

### Fluorescence imaging of mosquito midguts

Mosquitoes in No Abx, AbxA, and AbxPA groups that ingested ZIKV-spiked bloodmeals were imaged using immunofluorescence microscopy to examine midgut structure and to quantify the number of ZIKV-infected midgut epithelial cells. Legless and wingless mosquito bodies or dissected midguts from mosquitoes at 2 or 3 dpf were placed in 10% formalin in PBS for 24 hours, then prepared for various image-based assessments as described below. Mosquitoes were imaged at the UC Davis Advanced Imaging Facility using a Leica TCS SP8 STED 3X inverted microscope.

*Midgut structure*:Mosquito bodies were washed once in PBS, submerged in PBS with 30% sucrose overnight, and then molded and frozen at-80°C in Tissue Plus OCT medium (Thermo Fisher Scientific, Emeryville, CA). Tissue molds were cut into 10 µm slices at the median sagittal plane (at a depth where the blood bolus reaches maximum diameter, determined with test samples) using a Leica CM1860 Cryostat Microtome (Leica Biosystems, Deer Park, IL, USA) and embedded onto charged microscope slides. A single drop of ProLong Gold Antifade Mountant with DAPI (Thermo Fisher Scientific, Emeryville, CA) was used to mount each slide. *ZIKV infection of midguts:* Dissected midguts in 10% formalin were washed twice in PBS. Tissues were submerged in a blocking solution consisting of 1X PBS, 0.2% Triton X-100 (PBS+T) (Bio-Rad Laboratories, Hercules, CA), 1% bovine serum albumin (BSA, Sigma-Aldrich, St. Louis, MO), and 10% normal goat serum (Lampire Biological Laboratories, Pipersville, PA) for 10 minutes before staining with a primary monoclonal antibody that reacts with multiple flaviviruses (Flavivirus treatment antigen antibody D1-4G2-4-15, Novus Biologicals, Littleton, CO) at a 1:1000 dilution in blocking solution at 4°C overnight. Midguts were washed five times via direct tube transfer in PBS+T before staining with the secondary antibody (Goat anti-mouse Alexa Fluor 594, Abcam, Cambridge, United Kingdom) at a 1:1000 dilution in blocking solution for 1 hour at room temperature. Midguts were washed 5X in PBS+T before mounting onto microscope slides with ProLong Gold Antifade Mountant with DAPI. ZIKV detection and quantification were done by manually counting distinct red patches, using blue DAPI as a reference for individual cells across three slices of a single midgut, and averaging the values. FITC, DAPI, and Alexa Fluor 594 were each visualized (and FITC was quantified) in their respective channels before merging.

### Histologic analyses

Histological images at 20X and 40X magnification were used to visualize mosquito anatomy and to validate that sample preparation for immunofluorescence microscopy did not mechanically disrupt the midgut, which could affect interpretations of midgut integrity. Blood-engorged mosquitoes with their legs and wings removed were placed in 10% formalin in PBS for 24 h, then encased within tissue cassettes (Thermo Fisher Scientific, Emeryville, CA) submerged in 70% ethanol. Samples were submitted to the UC Davis School of Veterinary Medicine Anatomic Pathology Service, where whole mosquitoes were paraffin-embedded, cut into 5 µm slices, and stained with hematoxylin and eosin (H&E). Slides were visualized at 20X and 40X magnification under a Zeiss Axio Vert A1 inverted light microscope connected to an Axiocam 208 color camera (Carl Zeiss, Jena, Germany). Although we evaluated sections from 5 individual mosquitoes each for No Abx, AbxA, and AbxPA, groups, we used histology readouts as a qualitative approach to evaluate gross midgut anatomy. Anatomical structures were identified by an expert in mosquito anatomy. Representative images for each of the three groups are shown.

### Blood digestion analyses

Quantification of hemoglobin in bloodfed mosquitoes with a hemoglobin colorimetric detection kit (Thermo Fisher Scientific, Emeryville, CA) was used to measure blood digestion. The blood only and ZIKV-spiked bloodmeals each included 3 ml of blood. An additional 150 ul ZIKV was added to the ZIKV-spiked bloodmeal, such that the volume of virus was 5% of the total bloodmeal volume. At 0, 1, 2, or 3 dpf, 10-12 individual mosquitoes each in No Abx, AbxA, and AbxPA, groups exposed to blood only or ZIKV-spiked blood with their legs/wings removed were homogenized in 300 µL hemoglobin diluent, 100 µL of which was used for the assay, per manufacturer’s instructions. Absorbance at 570 nm was measured from each sample once in 96-well plates using a SpectraMax iD5 Multimode Microplate Reader (Molecular Devices, San Jose, CA). Absorbance values were converted to hemoglobin concentrations via standard curves generated from hemoglobin standards and serial dilutions of fresh sheep blood used for feeding. Output absorbance values were normalized to homogenized, non-bloodfed mosquito negative controls, and units were converted to µg per mosquito body. Before conducting the assay, we validated and optimized this kit for bloodfed mosquitoes, including ensuring that heat incubation of mosquitoes at 60°C for 10 minutes to destroy ZIKV infectivity did not affect hemoglobin quantification (**Figure S1**).**RNA sequencing and bioinformatics** Mosquito messenger RNA was sequenced to determine differences in gene expression associated with antibiotic treatments after ZIKV exposure. Total RNA was extracted from adult female mosquitoes in triplicate pools of 5 individuals each for No Abx, AbxA, and AbxPA groups using the Quick-RNA Tissue/Insect Microprep kit (Zymo Research, Irvine, CA, USA). RNA extracts were eluted into 20 µL elution buffer and quantified using a Qubit fluorometer RNA high-sensitivity (HS) assay (Thermo Fisher Scientific, Emeryville, CA). Approximately 1 µg of total RNA per sample was submitted to Genewiz Inc. from Azenta Life Sciences (Chelmsford, MA, USA), where poly-A selection, library preparation, and paired-end Illumina HiSeq 2×150 bp were performed. Raw FASTQ data files were received and treated by filtering and trimming reads with *cutadapt v3.2* (40) using default parameters (Quality cutoff >= 30, minimum length of 100 bp). Next, *FastQC* (41) was used to visualize quality scores. Processed reads were then aligned to version AaegL5.2 of an *Ae. aegypti* transcriptome obtained from VectorBase (42) using *bowtie2 v2.4.3* (43). Aligned reads were imported into R and analyzed with the package Differential Expression Analysis using *DESeq2 v1.28.1* (44). Default parameters were used to determine the differential abundance of each gene using an input experimental design of [antibiotic treatment + bloodmeal status + interaction]. Genes with a significance (adjusted P value) below 0.05 are reported. Significant genes were mapped to biological processes using the Gene Ontology program *topGO* (45).

### Statistical analyses

The 16S:RPS17 ratios to verify antibiotic efficacy from qPCR were compared using a Kruskal-Wallis test with multiple comparisons for adult mosquito treatments and across life stages. Differences in ZIKV infection, dissemination, and transmission rates between groups were determined using Fisher’s exact tests. Differences in ZIKV RNA levels in bodies, legs/wings, and saliva across groups were assessed by Kolmogorov-Smirnov tests. Differences in ASV and Evenness were evaluated using unpaired t tests. Differences in hemoglobin levels across groups were evaluated using Kruskal-Wallis tests with repeated measures. Relative fluorescence values and numbers of nuclei were averaged in triplicate and tested for significance by one-way ANOVA tests. Differences in numbers of ZIKV-infected midgut epithelial cells were assessed using Fisher’s exact tests. For RNA-Seq, statistical tests built into the *DESeq2* package were used. In short, each sample was normalized by library size, and the geometric mean for each gene was calculated. Negative binomial generalized linear models (glm) were used for each gene to determine differential expression across antibiotic-treated groups. The Wald test was used for statistical significance, with correction for the false discovery rate (FDR) set to 0.05. For principal components analyses (PCA), variance stabilized transformation was implemented before plotting. All statistical analyses, except RNA-Seq, were performed using GraphPad PRISM 9.0.2 (GraphPad Software, San Diego, CA). RNA-Seq analyses were conducted in R *v3.6* (46). P-values of less than 0.05 were considered statistically significant. For gene ontology (GO), Fisher’s exact tests were used on GO terms and corrected for the false discovery rate.

**Data availability.** Raw sequencing data are available from the NCBI Sequence Read Archive under BioProject entry PRJNA818687. Scripts for the RNA-Seq analysis are available on Github (DOI:10.5281/zenodo.7259822).

## Results

### Reduced *Ae. aegypti* susceptibility to ZIKV conferred by microbial exposure is life stage-dependent

In our previous study, we observed that adult *Ae. aegypti* reared in environmental water as larvae were less competent ZIKV vectors, assessed as lower infection and transmission rates, compared to *Ae. aegypti* reared in laboratory tap water (15). In this study, to understand whether reduced ZIKV vector competence is conferred transstadially from larvae to pupae to adults, we first determined whether exposure to microbes throughout life (i.e. at larval, pupal, and adult stages) is required to reduce ZIKV susceptibility. *Ae. aegypti* larvae were reared in laboratory water spiked with microbes that were collected from water in cemeteries that we characterized in a previous study (15) which contain >1000 bacterial species from the phyla *Bacteroides*, *Firmicutes*, and *Proteobacteria*. Control mosquitoes were not provided antibiotics (No Abx) at any life stage. To assess the effects of microbe modulation at pupal and adult life stages, cohorts of **p**upae and the **a**dults they developed into were administered an antibiotic (Abx) cocktail (AbxPA) that was provided to pupae in water and orally in sucrose to adults from eclosion to 3 days of age (**Figure 1A**). To study the effects of microbe modulation on adult mosquitoes that were not antibiotic treated at earlier life stages, we also included cohorts where **a**dults were provided antibiotics (AbxA) orally for 3-5 days post-eclosion. Antibiotics significantly reduced the number of prokaryotes in Abx treated adults as assessed by 16S:RPS17 ratios, our proxy for prokaryote levels (Kruskal-Wallis [K-W] with multiple comparisons, P = 0.002 between No Abx and AbxA [adj. P = 0.002] and No Abx and AbxPA [adj. P = 0.006]) (**Figure 1B**). Significantly higher 16S:RPS17 ratios were also observed in No Abx L4 larvae and pupae relative to adult mosquitoes (K-W multiple comparisons, P = <0.0001 between No Abx and larva [adj. P = 0.0012] and No Abx and pupa [adj. P = 0.0079]). This is consistent with ours (15) and others’ (7,13) observations that microbial abundance decreases during mosquito development. Mosquitoes were orally presented with ZIKV-spiked bloodmeals for vector competence assessments. AbxPA showed significantly increased ZIKV infection rates compared to No Abx (Fisher’s exact test, P < 0.0001) while AbxA and No Abx infection rates were not different (Fisher’s exact test:P = >0.05). Dissemination rates were not significantly different across treatments (Fisher’s exact test:P = >0.05). Significantly more AbxA mosquitoes transmitted compared to No Abx or AbxPA (Fisher’s exact test:P < 0.002) (**Figure 1C**). Levels of ZIKV RNA in saliva were higher in AbxPA compared to AbxA (Kolmogorov-Smirnov (KS) test, P = 0.009), but not higher than No Abx (KS test, P = 0.71) (**Figure 1D**). Collectively, these results show that microbe disruption across *Ae. aegypti* life stages increases rates of ZIKV infection (AbxPA), transmission (AbxA) and the magnitude of ZIKV transmitted in saliva (AbxPA).

**Figure 1.**
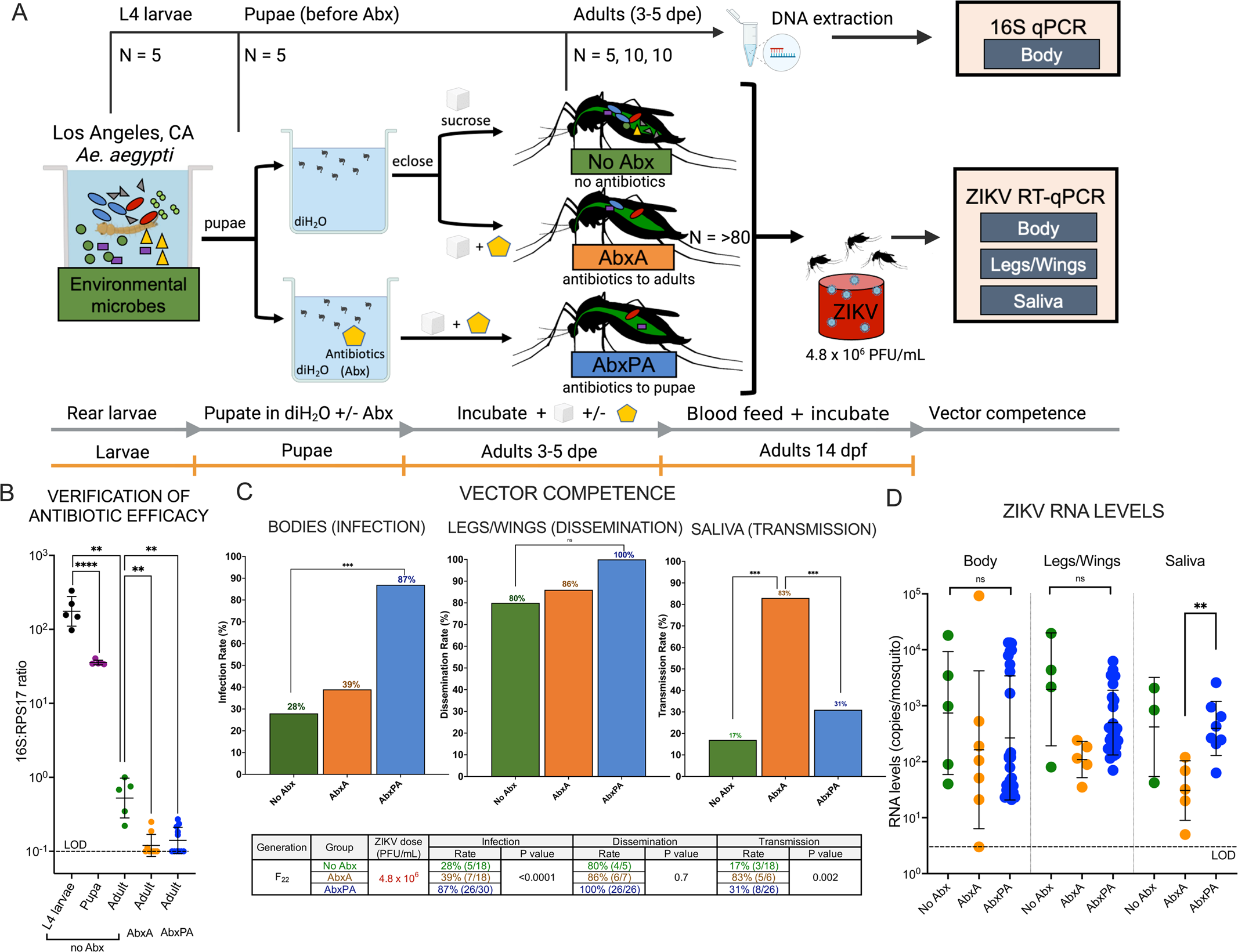
Microbial disruption using antibiotic treatment of *Ae. aegypti* pupae and adults from Los Angeles, California, increases ZIKV susceptibility. A) Experimental design. Microbes from cemetery headstones were added to tap water during rearing of *Ae. aegypti* larvae from Los Angeles (LA), CA. Pupae in the antibiotic (Abx) groups were treated by adding an antibiotic cocktail to pupal water and orally in sugar to adults (AbxPA). Antibiotics were delivered orally in sugar to adult Abx (AbxA) groups. No antibiotics (No Abx) groups received no antibiotics at any life stage. Mosquitoes from each group and some life stages were sequenced to verify the efficacy of antibiotic treatment. Adult mosquitoes 3-5 days of age were presented with ZIKV spiked into bloodmeals, and ZIKV vector competence was assessed by measuring ZIKV RNA levels in bodies to determine infection, legs/wings of individuals with infected bodies to assess dissemination, and in saliva of individuals with infected legs/wing to measure transmission, which are presented as rates in cohorts and mean levels from individuals. **B) Antibiotic treatment reduces bacterial DNA in mosquitoes.** Bacteria was quantified in mosquitoes by 16S qPCR. Each point represents a single mosquito (N larvae, pupae, No Abx = 5; N AbxA, AbxPA = 10). 16S copies were normalized to the *Ae. aegypti* RPS17 gene. The limit of detection (LOD) was a 0.1 16S:RPS17 ratio; below this value, 16S was not detected. Statistical significance across all groups was determined by a Kruskal-Wallis test with multiple comparisons. **C) ZIKV vector competence in microbe-modulated mosquitoes.** Bar graphs show ZIKV infection, dissemination, and transmission rates from cohorts as determined by ZIKV RNA RT-qPCR. The summary table shows the same data in the bar graphs. Rates are calculated as the number of bodies, legs/wings, or expectorates that yielded at detectable ZIKV RNA, divided by the total number of individuals that ingested blood within a cohort, were infected, or developed disseminated infections, respectively. Statistical significance across all treatments was determined by Fisher’s exact tests. Vector competence experiments were conducted once where mosquitoes in different cohorts were presented the same bloodmeal. **D) ZIKV RNA levels in mosquito tissues after ingestion of ZIKV-spiked bloodmeals**. Each dot shows the mean value for an individual mosquito based on triplicate measurements. The average LOD across all experiments was 3 copies per mosquito and is represented as the dotted line. Mosquitoes without detectable ZIKV RNA are not shown. Kolmogorov-Smirnov tests were used to compare mean RNA levels. For B and D geometric means are shown and error bars show standard deviations. *Denotes P < 0.05 **denotes P < 0.01, *** denotes P < 0.001, ns is not significant at P<0.05.

### Antibiotics do not directly impact midgut permeability or ZIKV vector competence

Next, we performed experiments to understand whether increased ZIKV vector competence in *Ae. aegypti* is due to direct effects of antibiotics on mosquito physiology or the dysbiotic microbial environment. We compared ZIKV vector competence and midgut integrity in mosquitoes treated with antibiotics (Conventional + Abx) or not antibiotic treated (Conventional), as well as in gnotobiotic mosquitoes (Gnotobiotic) (**Figure 2A**). For the antibiotic treated group, antibiotics were administered to both larval water and adults in sugar water using the same design as for the AbxPA cohorts. Gnotobiotic mosquitoes have a known microbiome compared to mosquitoes reared in environmental water, and, unlike axenic mosquitoes that do not possess any microbes, do not suffer from the lack of microbiome-derived essential nutrients that can impact fitness (47). Gnotobiotic *Ae. aegypti* were generated by inoculating larval water with an *Elizabethkingia spp.* culture isolated from colonized *Ae. aegypti* from LA. *Elizabethkingia spp*. was selected since it is ubiquitous in the environment (47) and represents a dominant bacterial sequence in our larval and adult *Ae. aegypti* colonies (15). We verified the gnotobiotic status by sequencing (**Figure 2B**). Gnotobiotic mosquitoes showed significantly lower mean observed ASV and evenness (where lower evenness indicates dominance by fewer species) compared to Conventional or Conventional+Abx (p<0.05, unpaired t test). A mean of 180 ASVs were detected in Conventional mosquitoes, Conventional +Abx had 139, and Gnotobiotic had 47. Conventional mosquitoes reared with or without antibiotics had *Chryseobacteria* as the dominant ASV while in gnotobiotic mosquitoes, *Elizabethkingia* and *Asaia* comprised >95% of all observed ASVs (**Figure S2**). The co-dominance of our intended *Elizabethkingia* inoculum and *Asaia* spp., a commonly found bacterium in our mosquito colony that may be vertically transmitted (15), supports a successful gnotobiotic state. The three groups were presented with high dose ZIKV-spiked blood with and without fluorescent dextran (FITC-dextran). A subset of mosquitoes were evaluated within 1 hour of bloodfeeding for midgut permeability assays and the remaining bloodfed individuals were assessed for vector competence at 14 days. The purpose of adding FITC-dextran, which is a small molecule that can traverse cell membranes, was to evaluate whether antibiotics or a gnotobiotic environment influence midgut permeability. Infection, dissemination, and transmission rates were not statistically different across any group (Fisher’s exact test:P >0.09) (**Figure 2C**). Mean ZIKV RNA levels in ZIKV-positive bodies, legs/wings, and saliva were also not statistically different across groups (KS test, P >0.05) (**Figure 2D**). We visualized and quantified the relative fluorescent intensity of FITC-dextran inside versus outside the midgut (**Figure 2E**), where higher fluorescence inside versus outside is indicative of greater midgut integrity. Dextran permeability measurements showed no significant difference across groups in mean fluorescence inside versus outside of the midgut (KS test, P >0.05) (**Figure 2F**). These data indicate that antibiotics themselves or a gnotobiotic state do not perturb midgut permeability or affect ZIKV vector competence and rule out confounding effects of these treatments on the structural integrity of the mosquito midgut epithelium.

**Figure 2:**
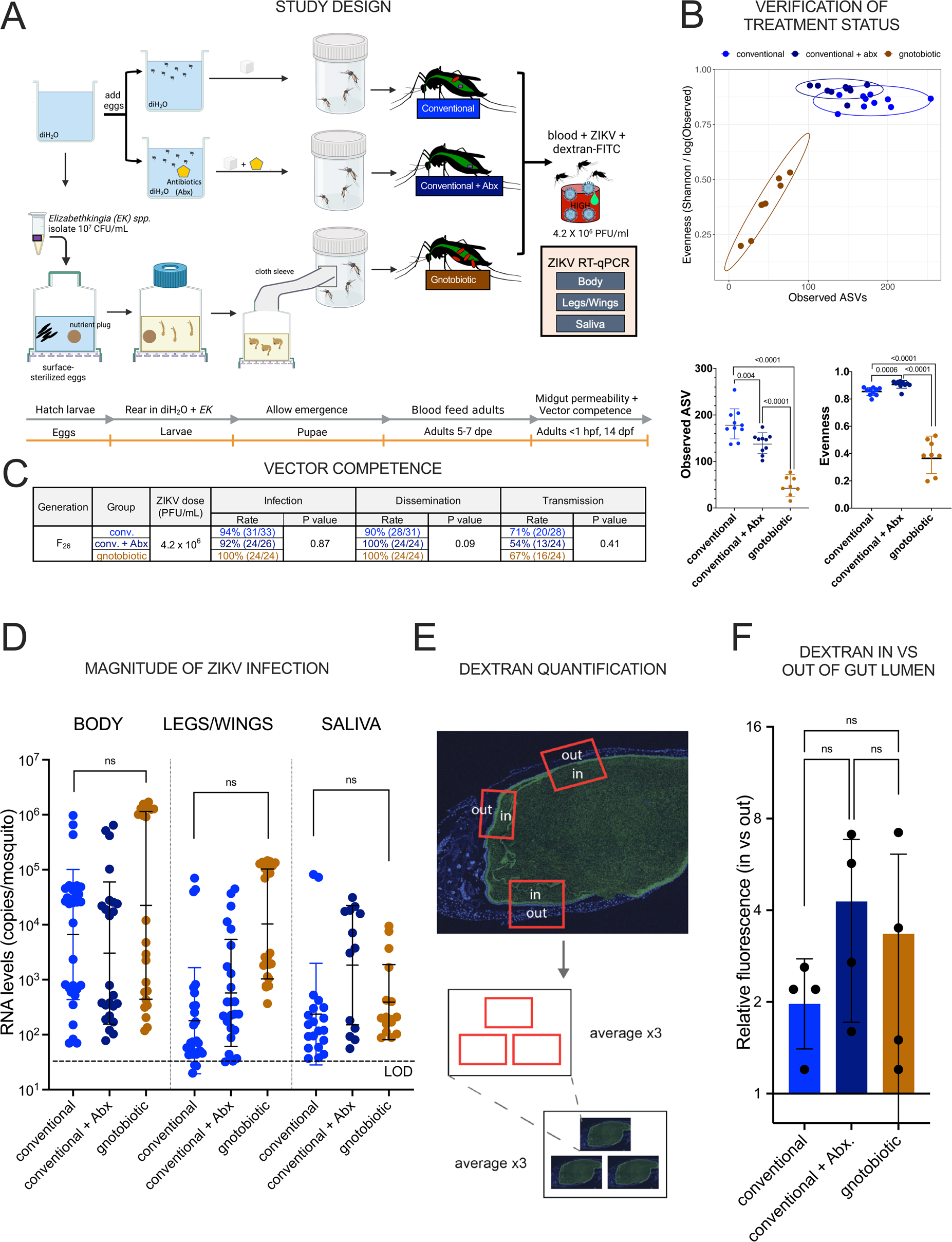
A) Experimental design. *Aedes aegypti* from LA, CA, were reared in laboratory tap water without any antibiotics (Conventional), with antibiotics (Conventional + Abx), where antibiotics were also provided orally to adults in sugar, or with *Elizabethkingia spp*. (Gnotobiotic) bacteria isolated from our *Ae. aegypti* colony. Mosquito eggs in the Gnotobiotic group were surface sterilized before hatching and reared in sterile flasks. Adult mosquitoes 3-5 days of age were presented with ZIKV spiked into bloodmeals, and ZIKV vector competence was assessed by measuring ZIKV RNA levels in bodies to determine infection, legs/wings of individuals with infected bodies to assess dissemination, and in saliva of individuals with infected legs/wing to measure transmission, which are presented as rates in cohorts and mean levels from individuals. The bloodmeals also contained 1 µg dextran-FITC (green drop) for midgut permeability assessments. **B) Verification of treatment status using 16S sequencing from adult mosquitoes in each treatment group.** The scatterplot shows the relationship between observed ASVs and bacterial species evenness. ASVs with unassigned taxonomy were included. Evenness is calculated by dividing Shannon diversity by the natural log of observed ASVs. Ellipses encompass the 95% confidence interval for samples within the group. Each dot on Observed ASV and Evenness graphs represents an individual mosquito (N=10 Conventional, N=10 Conventional + Abx, N=8 Gnotobiotic). P values reported were determined using unpaired t tests. **C) ZIKV vector competence** data comparing infection, dissemination, or transmission rates in Conventional, Conventional + Abx, and Gnotobiotic *Ae. aegypti* groups. Vector competence experiments were conducted once where mosquitoes in different cohorts ingested the same bloodmeal. Statistical evaluation across all treatments used Fisher’s exact tests. **D) ZIKV RNA levels in mosquito tissues after ingestion of ZIKV-spiked bloodmeals**. Each dot shows the mean value for an individual mosquito based on triplicate measurements. The average LOD across all experiments was 62 copies per mosquito and is represented as the dotted line. Mosquitoes without detectable ZIKV RNA are not shown. **E) Dextran fluorescence quantification approach** that was used to generate data in panel F. Fluorescence ratios were calculated in triplicate from randomly selected areas in a single midgut slice. Dextran permeation was calculated as the ratio of fluorescence intensity inside the gut lumen to fluorescence intensity outside the gut lumen for randomly selected regions of the midgut (red rectangles). **F) Mosquito midgut permeability assessed by relative dextran inside versus outside the midgut lumen**. Each symbol represents 9 measurements from one mosquito, N=4 mosquitoes per treatment. For C and E geometric means are shown, and error bars show standard deviations. Kolmogorov-Smirnov tests were used to compare mean RNA or fluorescence levels. ns is not significant at P<0.05.

### Microbe disruption increases *Aedes aegypti* midgut permeability

To understand how microbial exposure of larvae, pupae, and adults reduces susceptibility of *Ae. aegypti* to ZIKV, we applied the midgut permeability assays (**Figure 2E**) to evaluate ZIKV-exposed mosquitoes that were microbe-exposed or microbe-disrupted. Our rationale for focusing on the midgut is that it represents the first barrier to infection of *Ae. aegypti* by ZIKV after oral exposure. We asked whether higher infection rates in AbxPA compared to No Abx are explained by reduced midgut integrity resulting from microbe disruption. We compared fluorescent intensity across antibiotic-treated groups immediately after ingestion of a bloodmeal with ZIKV (**Figure 3A**). All mosquitoes had only bloodfed once, eliminating potential confounding effects of repeated bloodfeeding on midgut integrity (48) and were fully engorged; hemoglobin measurements (see below) also revealed no statistically significant differences in mean ingested bloodmeal volumes across groups. The relative dextran-FITC fluorescence inside versus outside of the midgut lumen for the AbxPA group was significantly lower than the No Abx group (P = 0.009), and the AbxA group trended lower (although not significantly) than the No Abx group (**Figure 3B**). To rule out physical rupture or other disruptions to the midgut during sample preparation that may have reduced structural integrity, we treated and stained midgut sections from a separate cohort of similarly treated mosquitoes with DAPI or H&E. At 10X, 20X, and 40X magnification, midgut epithelia across treatments did not appear structurally different (**Figures 3D**, **S1**). Together these data support a process where disruption of microbial colonization at pupal and adult *Ae. aegypti* life stages increase midgut cell permeability in adult mosquitoes after bloodmeal ingestion.

**Figure 3.**
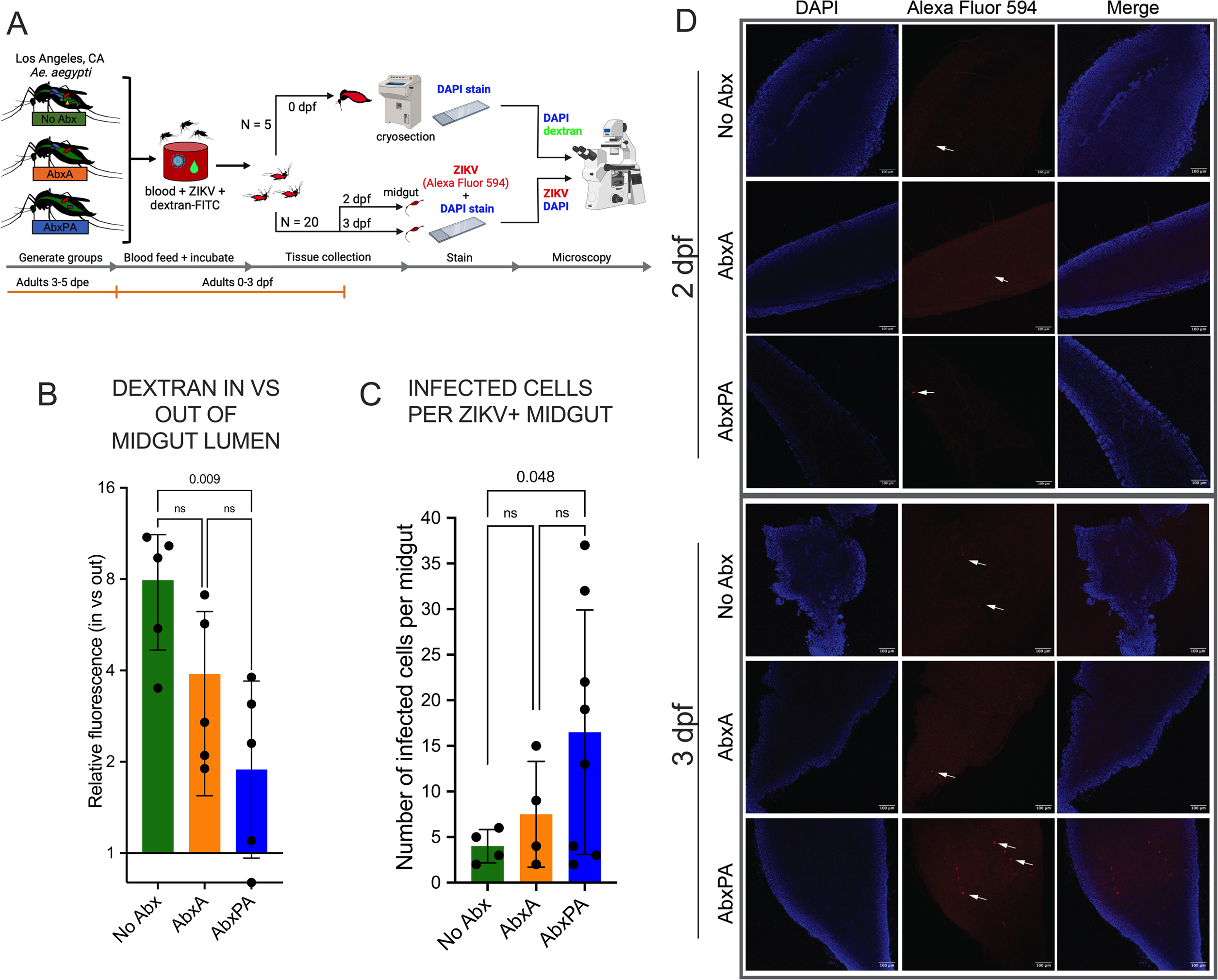
Microbial disruption increases *Aedes aegypti* midgut permeability. A) Schematic of bloodfeeding experiment and sample preparation for immunofluorescence microscopy. *Aedes aegypti* from LA, CA were used to generate three groups of mosquitoes:no antibiotics (No Abx), antibiotics as adults (AbxA), and antibiotics as larvae, pupae, and adults (AbxPA). Adult mosquitoes aged 3-5 days were presented with a bloodmeal containing 1 mg/mL dextran-FITC (green) and 3.3 x 10^6^ PFU/mL ZIKV. Whole mosquitoes were harvested within 1 hour of feeding at 0 days post-feed (dpf), cryosectioned, DAPI stained, and examined using fluorescence microscopy. Midguts from the same cohorts were harvested at 2 and 3 dpf, and ZIKV antigen and DAPI were stained before fluorescence microscopy. **B) Microbe disruption of pupae and adults increases midgut permeability, evidenced by reduced relative dextran inside versus outside the gut lumen in AbxPA compared to No Abx mosquitoes**. Dextran permeation was calculated as the ratio of fluorescence intensity inside the gut lumen to fluorescence intensity outside the gut lumen using the same approach as in Figure 2E. Each symbol represents 9 measurements from one mosquito. **C) Mosquitoes that were antibiotic treated as pupae and adults have more ZIKV-infected midgut epithelial cells**. The number of ZIKV-infected cells, as determined by ZIKV antigen staining, were counted for each ZIKV-positive midgut on 2 and 3 dpf. Data from both days is shown. Each symbol represents one mosquito. **D) Antibiotic treated ZIKV-positive midguts at 2 and 3 dpf.** Midguts examined under immunofluorescence at 10X magnification, at 2 and 3 dpf. Cell nuclei (blue) and ZIKV antigen (red) are shown with the combined (merged) channels. Images reflect representative observations for each antibiotic treatment. Arrows point to ZIKV antigen positive cells. P values in B and C use ANOVA with multiple comparisons and error bars show standard deviations. ns is not significant.

### Microbe disruption via antibiotic treatment increases ZIKV infection of the midgut epithelium

Next, we asked microbe disruption leads to increased midgut infection success, measured as higher numbers of ZIKV-infected midgut epithelial cells. We detected ZIKV in midguts by staining sections with a flavivirus binding antibody 2 or 3 dpf and counting the total number of cells in each midgut that were stained with fluorescent anti-ZIKV antibody (**Figure 3A**). The number of ZIKV-positive cells in midguts was significantly higher in the AbxPA group compared to the No Abx group (Brown-Forsythe ANOVA test with Dunnett’s multiple comparisons test, P = 0.048) (**Figure 3C**). These data support the role of microbes in larvae and pupae in maintaining the integrity of the mosquito midgut which associates with reduced midgut epithelium infection by ZIKV.

### Microbial disruption via antibiotic treatment impairs blood digestion

Since midgut integrity can affect physiology of blood digestion in addition to ZIKV susceptibility, we next assessed whether elevated midgut permeability stemming from microbe reduction by antibiotic treatment affects blood digestion. We employed a commercially available hemoglobin colorimetric assay as a proxy for blood protein digestion by quantifying the reduction of hemoglobin levels in blood-engorged mosquitoes on 1 and 2 dpf compared to immediately after feeding on 0 dpf. Before performing the assay, which is typically used for human blood, we first validated and optimized the approach for use with homogenized mosquitoes that ingested sheep bloodmeals. We observed that sheep blood absorbance varies linearly as a function of hemoglobin concentration, like the kit standard (**Figure S3A**) and that absorbance decreases reproducibly in serially diluted mosquitoes at both 560 and 580 nm wavelengths. The assay was unaffected by the 10-minute 60°C heat treatment of homogenized fed mosquitoes used to inactivate ZIKV infectivity in the samples (**Figure S3B**). Blood-engorged females were homogenized and assayed for hemoglobin immediately after bloodfeeding at 0 dpf to assess whether different groups ingested the same bloodmeal volume and to measure kinetics of bloodmeal digestion 1, 2, and 3 dpf (**Figure 4A**). Although hemoglobin levels that approximate bloodmeal size varied in individual mosquitoes, there was no significant difference in mean ingested hemoglobin levels on 0 dpf across antibiotic-treated groups regardless of whether the bloodmeal contained ZIKV, which shows that antibiotic treatment did not influence bloodmeal size (KW test with multiple comparisons, P = 0.522) (**Figure 4B,C**). After ingestion of blood without ZIKV, mean hemoglobin levels in the AbxPA group were significantly higher than in the No Abx group 2 dpf (KW test, P = 0.0063). By 3 dpf, mean hemoglobin levels in the AbxPA group were significantly higher than both AbxA and No Abx groups (KW test with multiple comparisons, adjusted P_AbxPA-No Abx_ = 0.0347, adjusted P_AbxPA-AbxA_ = 0.051) (**Figure 4B**). The same pattern was observed after ingestion of blood with ZIKV:AbxPA showed a significantly reduced rate of blood digestion compared to No Abx 1 dpf (KW test with multiple comparisons, adjusted P_AbxPA-No Abx_ = 0.0368, adjusted P_AbxPA-AbxA_ = 0.0035) and both No Abx and AbxA 2 dpf (KW test with multiple comparisons, adjusted P_AbxPA-No Abx_ = 0.0037, adjusted P_AbxPA-AbxA_ = 0.0027) (**Figure 4C**). Hemoglobin levels at 3 dpf for individuals that ingested blood with ZIKV was not assayed since uneven bloodfeeding rates across cohorts and the desire to maintain statistically robust group sizes led us to prioritize use these mosquitoes for infection and dissemination assessments. We also compared mean hemoglobin levels across treatments and days post-feed for mosquitoes that ingested blood with or without ZIKV. Although levels did not differ significantly after bloodmeal ingestion on 0 dpf, mean hemoglobin digestion was significantly faster in No Abx and AbxA on 1 dpf (KW test, P_No Abx_ = 0.0163, P_AbxA_ = 0.0009) but not AbxPA (KW test, P = 0.25) in ZIKV-exposed mosquitoes compared to mosquitoes that ingested blood without ZIKV (**Figure 4D**). To evaluate whether these observations extend to *Ae. aegypti* from a different genetic background than LA, CA, we performed the similar analyses with *Ae. aegypti* from Clovis, CA. Genome sequencing shows that *Ae. aegypti* from LA and Clovis are genetically distinct (49), and were probably derived from separate independent introductions into CA since 2013 when the species became established in the state. Using a similar design (**Figure S4A**), as for mosquitoes from LA, AbxA and AbxPA from Clovis showed significantly lower blood digestion relative to No Abx 2 dpf (KW test with multiple comparisons, adjusted P_AbxPA-No Abx_ = 0.0013, adjusted P_AbxPA-AbxA_ = 0.027) (**Figure S4B**). Clovis AbxPA exhibited a marginally but not statistically significantly higher infection rate (34%) compared to AbxA (26%) and No Abx (15%) groups (Chi-square [χ^2^] test, P = 0.106). The dissemination rates and mean RNA levels in ZIKV-positive Clovis mosquitoes were not different for any treatment (**Figure S4C**). These data show that disruption of larval and pupal microbes reduces bloodmeal digestion in two genetic lineages of *Ae. aegypti*, and that blood with ZIKV is digested more quickly than blood lacking virus.

**Figure 4.**
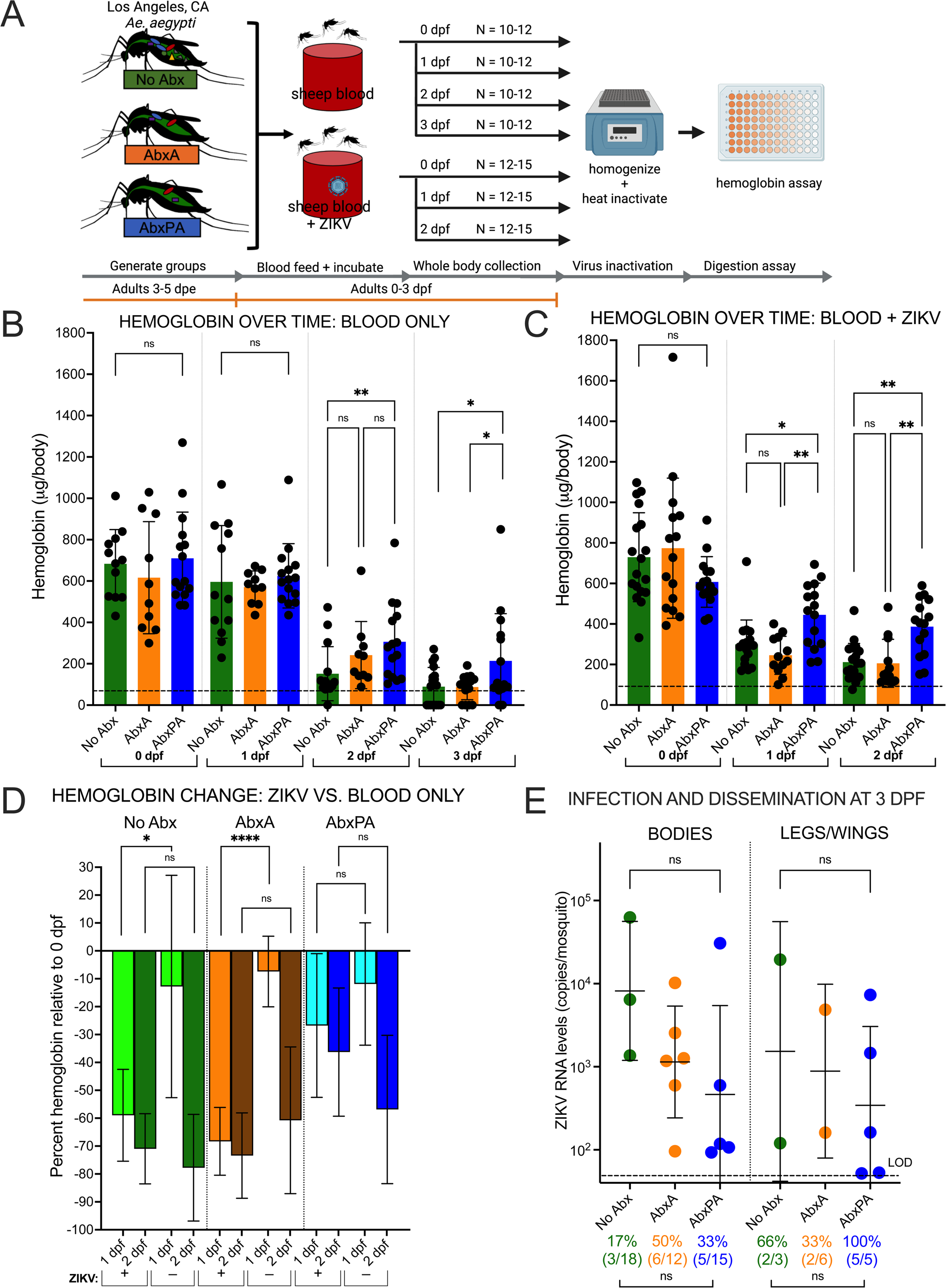
Bloodmeal digestion kinetics in antibiotic-treated microbe-disrupted *Ae. aegypti*. **A)** Experimental design for detection of hemoglobin in homogenized antibiotic treated mosquitoes that ingested blood with or without ZIKV. *Aedes aegypti* from LA, CA were reared as in Figure 1A to generate three groups of mosquitoes:no antibiotics (No Abx), antibiotics as adults (AbxA), and antibiotics as larvae, pupae, and adults (AbxPA). Adult mosquitoes aged 3-5 days were presented with a bloodmeal with or without 2.1 x 10^6^ PFU/mL ZIKV. On day 0 (within 1 hour of feeding) and at 1, 2 and 3 days dpf a subset of 10-15 mosquitoes were homogenized and heat treated to destroy ZIKV infectivity for bloodmeal volume detection using a hemoglobin assay. Infection and dissemination were assessed 3 dpf. **Hemoglobin levels in *Ae. aegypti* as a proxy for bloodmeal digestion in groups that were (B) bloodfed or (C) bloodfed with ZIKV.** The number of *Ae. aegypti* that ingested blood that were assayed per day is:No Abx = 12, AbxA = 10, AbxPA 15. The number of *Ae. aegypti* that ingested blood spiked with ZIKV that were assayed per day is:No Abx = 18, AbxA = 12, AbxPA = 15. **D) Hemoglobin levels in bloodfed versus ZIKV exposed *Ae. aegypti*.** Mosquitoes that ingested ZIKV that were not treated with antibiotics or treated only as adults show increased blood digestion compared to mosquitoes treated with antibiotics as pupae and adults. The bar graphs show the percent reduction in hemoglobin at 1 and 2 dpf compared to the 0 dpf, calculated for all Abx groups and between ZIKV exposed and bloodfed only groups. **E) Microbial disruption of pupae and adults does not impact ZIKV dissemination rates 3 dpf.** Legs and wings of individual bloodfed females from the same cohort were assayed by ZIKV RT-qPCR at three dpf. Each symbol in B, C and E shows the mean measurement for 1 mosquito. Bars show mean and error bars indicate the standard deviation. * Denotes P < 0.05; ** denotes P < 0.01; **** denotes P<0.0001, ns is not significantly different.

### Reduced bloodmeal digestion does not influence ZIKV dissemination from the *Ae. aegypti* midgut

We next hypothesized that, in addition to disrupting midgut cell integrity, antibiotic mediated reduction of microbes in mosquitoes and resulting reduced bloodmeal digestion provides the virus more time to bind and enter midgut epithelial cells, manifest as increased ZIKV infection and dissemination very early (3 dpf) after bloodfeeding. However, the rates and magnitude of ZIKV RNA in bodies and legs/wings in antibiotic-treated groups was not different from the mosquitoes that were not antibiotic treated that ingested ZIKV-spiked blood 3 days earlier (Fisher’s exact test, P > 0.05) (**Figure 4E**). Together with the midgut cell infectivity data, these results suggest that increased ZIKV midgut cell infection observed in antibiotic treated pupae and adults establishes after 3 dpf and that augmented rates of infection in microbe-disrupted mosquitoes do not augment early dissemination.

### Hemolytic bacteria do not augment bloodmeal digestion in *Ae. aegypti* that were not microbe modulated via antibiotic treatment

To determine whether bacteria with hemolytic activity contribute to enhanced bloodmeal digestion in the No Abx groups, bacteria from 5 No Abx treated adult mosquitoes were cultured on LB and R2A agar and then Sanger sequenced. Although different species of bacteria were detected, none showed hemolytic activity (**Table S1**), suggesting that bloodmeal digestion is not directly mediated by bacteria in the mosquito midgut in the *Ae. aegypti* studied with the assays used here. Together, these data show that microbes enhance the ability of *Ae. aegypti* to digest blood, likely via a mechanism not tied directly to bacteria mediated hemolysis since the culturable bacteria recovered from mosquitoes that were not antibiotic treated were not hemolytic.

### Blood ingestion and ZIKV exposure influence the *Ae. aegypti* transcriptome

Absent increased dissemination from the midgut, to identify gene expression differences associated with increased ZIKV infection in AbxPA, we employed RNA-Seq analyses on pooled adult female *Ae. aegypti* from LA before bloodfeeding and at 2 dpf (**Figure 5A**). We also analyzed females that ingested blood with our without ZIKV. On average, 94.5% (min:92.7, max:95.7%) of reads from all samples aligned at least once to the Liverpool *Ae. aegypti* transcriptome. Mosquitoes that had not bloodfed clustered separately from those that ingested blood with or without ZIKV (**Figure 5B**, **Figure S5A**). Bloodmeal status correlated more strongly with variance (accounting for 84%, P<0.001) in the overall transcriptomic profile compared to antibiotic treatment (<5%, P = 0.83). ZIKV in the bloodmeal also yielded differentially expressed genes (DEGs) for each antibiotic treated group. Prior to blood with or without ZIKV, 20 significant DEGs were identified between AbxPA and No Abx (**Table S2**):the most downregulated in AbxPA relative to No Abx was uncharacterized and other downregulated genes in AbxPA included membrane components and ATP binding proteins. After bloodfeeding, over a third of genes in the *Ae. aegypti* genome were differentially expressed, consistent with other RNA-Seq studies (50). At 2 dpf, 7846 genes in No Abx, 9356 genes in AbxA, and 7846 genes in AbxPA were differentially expressed compared to sugar-fed only mosquitoes. Among them, 6676 genes were shared among No Abx, AbxA, and AbxPA, representing an overwhelming majority of the total DEGs (**Figure S5B**). Between bloodfed without and with ZIKV, DEG differences were more apparent:AbxA had 1974 DEG, and No Abx had 1258, and AbxPA had 62 (**Figure 5C**, only DEGs that differed significantly across antibiotic treatments are shown [adjusted P < 0.05 by Wald test]). Most shared DEGs were between AbxA and No Abx groups; AbxPA expressed a unique set of DEGs. Assignment of Kyoto Encyclopedia of Genes and Genomes (KEGG) pathway identity (IDs) to DEGs identified many unknown genes. Hypothetical protein products with no homology to genes with functional annotations were common (No Abx:526, AbxA:792, AbxPA:29), as were genes involved in metabolic processes (No Abx:371, AbxA:599, AbxPA:25) and thermogenesis (No Abx:277, AbxA:452, AbxPA:9) (**Figure 5D**). In ZIKV bloodfed mosquitoes, AbxPA reduced differential gene expression more than AbxA or NoAbx. Taken together, these data suggest that bloodfeeding has a larger effect than microbial exposure on *Ae. aegypti* gene expression and implicate microbe-mediated effects on factors other than changes in transcription as modifiers of ZIKV vector competence.

**Figure 5.**
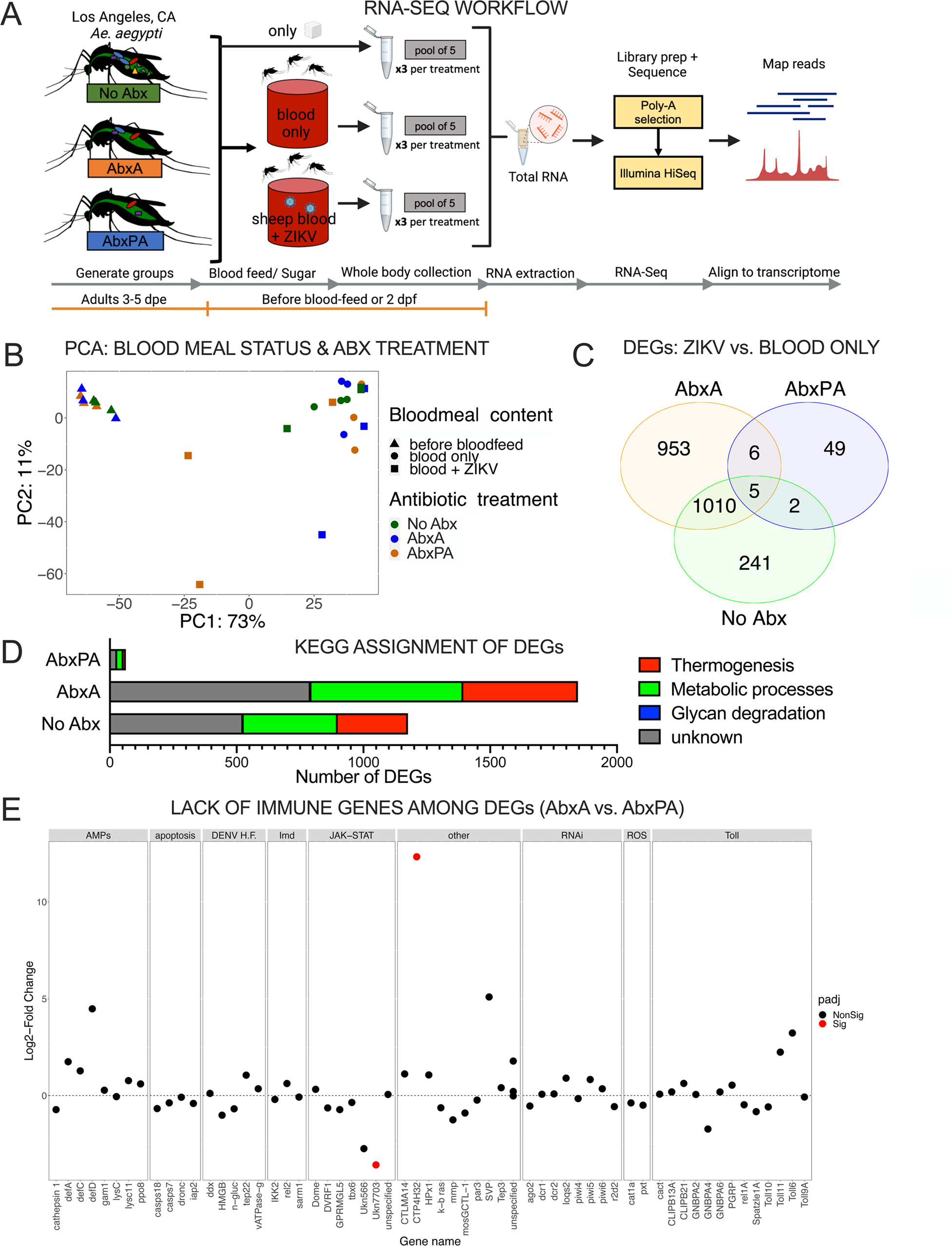
Differentially expressed genes between antibiotic-treated adult *Ae. aegypti* females that had or had not taken a bloodmeal. A) Experimental design. For each antibiotic treatment and bloodfeeding state, triplicate pools of 5 whole, legless mosquito bodies were RNA-extracted, pre-processed, and sequenced. **B) Transcriptome profile differs more by bloodmeal status than by antibiotic treatment.** PCA plot after variance stabilized transformation. Shapes indicate the bloodmeal status and colors represent antibiotic treatments. Separate PCAs for individual variables, are shown in **Figure S5**. C) ZIKV-responsive DEGs are shared between *Ae. aegypti* that were not antibiotic-treated or antibiotic-treated as adults. Venn diagram of DEGs in non-bloodfed and bloodfed *Ae. aegypti* for each treatment. **D) DEGs correspond to metabolic processes, thermogenesis, and unknown genes.** KEGG Pathway ID assignment of DEGs in panel C. E) DEGs between *Ae. aegypti* mosquitoes treated with antibiotics as pupae and adults compared to adults only and between mosquitoes that ingested ZIKV in blood or blood only do not correspond to known immune response genes. Relative expression of select immune-related genes from literature (51–66). Expression is higher in AbxA if the log_2_-fold change is positive, while expression is higher in AbxPA if log_2_-fold change is negative. Points are considered significant (red) if the adjusted P value is < 0.01. Comparisons of the same genes between AbxA or AbxPA and the control treatment, No Abx, are shown in **Figure S6**.

Provided that microbes mediate immunomodulatory effects in mosquitoes (13,67,68), we theorized their presence could influence ZIKV susceptibility by modifying the expression of immune response pathways where some respond to both bacteria and viruses. We identified immunity-related DEGs across antibiotic-treated mosquitoes 2 dpf in the RNA-Seq data. Transcripts representing components of various immune pathways were accessioned from existing literature and used to filter the dataset before analyses of DEG profiles. Representative genes examined include regulators from Toll (51–53), RNAi (54–57), JAK-STAT (52,58,59), and Imd pathways (60,69), as well as general immune and immune-adjacent genes such as hormones (61–63,70), antimicrobial peptides (AMPs) (59,64), reactive oxygen species (ROS) (65,66), and known DENV host factors (59,64). There were few DEGs demonstrating significant changes in transcript abundance in immune pathways between AbxA and AbxPA, No Abx and AbxA, or No Abx and AbxPA groups (**Figures 5E, S5**). Comparison of immune genes transcript abundance differences between No Abx and AbxA and between No Abx and AbxPA revealed several JAK-STAT pathway genes (**Figure S6**). These data suggest that gene expression changes after microbe disruption via antibiotic treatment of *Ae. aegypti* that show different susceptibility to ZIKV infection are not related to known immune response pathways.

### Gene ontology assignments reveal the role of microbiota in stimulating expression of protein digestion-associated genes in *Ae. aegypti* orally exposed to ZIKV

Given that RNA-Seq revealed no major differences in expression of immune response genes after microbe disruption, we next performed Gene Ontology (GO) analyses to predict enriched functions in non-immune associated DEGs. We identified DEGs displaying significant changes in abundance between antibiotic treated mosquitoes that ingested ZIKV in blood or that ingested blood only. The AbxA and No Abx groups that ingested ZIKV in blood showed increased expression of genes enriched in general translation processes (AbxA:125, adjusted P < 10^-10^; No Abx:127, adjusted P < 10^-20^) and proteolysis (AbxA:84, adjusted P = 0.0006; No Abx:56, adjusted P = 0.015). The AbxPA group showed few upregulated genes resulting in a lack of enriched functions (**Figure 6A**). Conversely, enrichments in genes downregulated in mosquitoes that ingested ZIKV in blood versus mosquitoes that ingested blood showed reduced transcript abundance for genes associated with DNA metabolism (8) and redox processes (4) in the ZIKV exposed AbxPA mosquitoes. AbxA and No Abx were depleted in genes for DNA replication (N=48 and 34, respectively) and DNA repair (N=18 and 2, respectively). All treatment groups that ingested blood with ZIKV compared to blood only exhibited downregulation of carbohydrate metabolism genes (AbxPA:N=8, adjusted P = 0.006; AbxA:N=6, adjusted P = 0.0002; No Abx:N=16, adjusted P = 0.017) (**Figure 6B**). These data suggest that microbiota during early ZIKV infection promote expression of genes associated with proteolysis and translation while downregulating genes relating to DNA synthesis. The magnitude of change in DEGs in antibiotic treated mosquitoes that ingested ZIKV in blood or blood only were also compared. Gene AAEL008117, the most depleted transcript in AbxA and No Abx groups (∼20 log_2_-fold reduced), corresponds to nompC, a mechanosensitive ion channel that senses balance and touch in *Drosophila melanogaster* (71). The most common DEGs that changed in response to ZIKV exposure matched to proteolytic and PM-associated enzymes, including endopeptidases, collagenases, and dehydrogenases (72). Multiple serine-type endopeptidases were differentially expressed depending on antibiotic treatment (**Figure 6C**). In addition, some vitellogenesis associated proteins were more enriched in antibiotic treated mosquitoes compared to the No Abx group. In AbxPA, the gene demonstrating the biggest reduction in transcript abundance corresponds to a m1 zinc metalloprotease, showed increased expression in the No Abx group. Overall, GO analyses revealed *Ae. aegypti* with disrupted microbiota are deficient in genes that influence blood protein metabolism.

**Figure 6.**
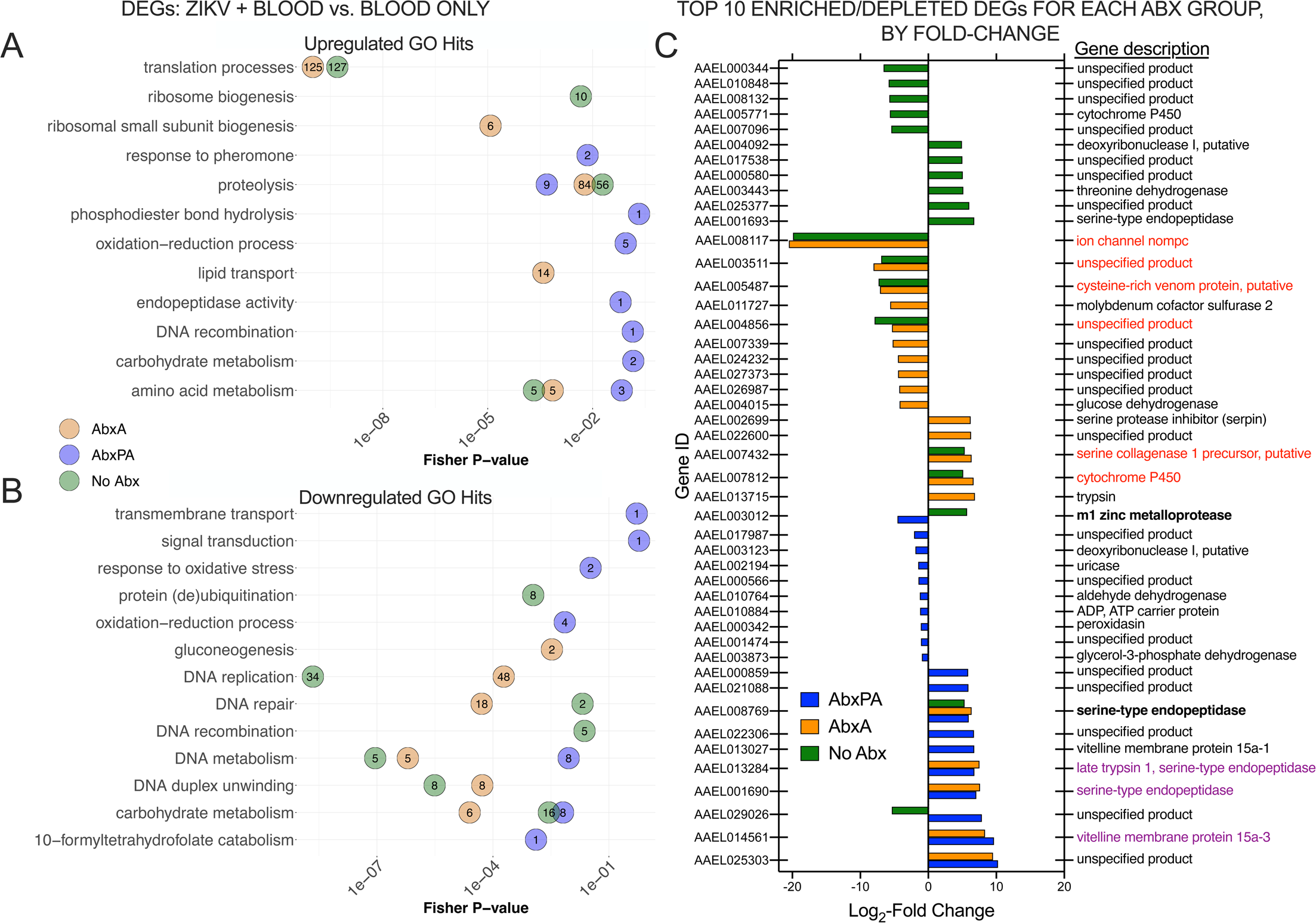
Gene Ontology assignment of DEGs and identity of most enriched/depleted genes in *Ae. aegypti* that were antibiotic treated and that ingested ZIKV in bloodmeals or blood only. **A) Upregulated GO hits for mosquitoes treated with antibiotics as adults or not antibiotic treated map to general translational processes and proteolysis.** Mosquitoes treated with antibiotics as pupae and adults demonstrate a reduced complement of proteolysis associated genes, and **B) GO enrichments in genes showing reduced transcript abundance for all antibiotic treatments primarily map to DNA synthesis and repair mechanisms.** GO terms for the DEGs (determined from Figures 6C-D) and their descriptions on the y-axis, and statistical significance (P-values, Fisher’s exact test) of the GO term on the x-axis, with the exact number of DEGs within the colored circles. DEGs farther to the left (lower P-values) are more significant. **C) ZIKV-responsive DEGs demonstrating the highest magnitude fold change are distinct between AbxPA and No Abx treatment groups.** The top 10 upregulated and top 10 downregulated DEGs, by magnitude log_2_-fold change for each antibiotic treatment. A positive log_2_-fold change refers to more transcripts after a ZIKV-spiked bloodmeal, while a negative value refers to fewer transcripts. The right y-axis is the gene description from annotations retrieved from transcriptome metadata as well as VectorBase (42). Red text indicates the top genes shared between No Abx and AbxA. Purple text indicates the top genes shared between AbxPA and AbxA.

## Discussion

This study shows that microbial exposure confers resistance to ZIKV infection of *Ae. aegypti* and that microbes enhance bloodmeal digestion and midgut integrity and reduce midgut cell infection. Reduced microbial colonization during both pupal and adult *Ae. aegypti* life stages produced the largest increases in ZIKV susceptibility. These effects were not due to antibiotic treatment itself and our studies with gnotobiotic mosquitoes show that exposure to more than *Elizabethkingia spp.* is required to reduce ZIKV susceptibility. The lack of modified ZIKV susceptibility in microbe-reduced Clovis *Ae. aegypti* contrasts with our observations from Los Angeles mosquitoes and highlights the need for further work to understand whether microbes have differential impacts on ZIKV competence in *Aedes* mosquitoes from different regions, which is supported by a recent study showing mosquito genotype effects in Senegalese *Ae. aegypti* (18).

Augmented rates of ZIKV infection but not dissemination in *Ae. aegypti* after microbe reduction via antibiotic treatment suggest that microbes act at the midgut infection barrier. Augmented transmission by AbxA but not AbxPA could be explained by a stochastic effect due to the smaller group size of AbxA versus AbxPA or another mechanism we do not yet understand. Hematophagous feeding introduces a unique problem in that engorgement immensely distends the midgut. Midgut distention can result in breaches in the integrity of the epithelial barrier, presenting an opportunity for a virus ingested with blood to bypass infection of the epithelium. Prior studies detected bloodmeal-induced perforations in the midgut basal lamina, which do not fully recover after digestion, supporting a process where successive bloodmeals increase rates of ZIKV dissemination (48). Considering that we observed a reduced magnitude of and rate of bloodmeal digestion in antibiotic-treated mosquitoes as pupae and adults, we envision a process where microbes accelerate digestion, decreasing the window of time the virus has to escape the bloodmeal and access the midgut epithelium. We did not observe gross structural differences in the midgut epithelium with DAPI or H&E staining, so we theorize that microbes do not impact midgut permeability by physically modifying the midgut epithelium, although electron microscopy studies could yield more insight into structural changes. We acknowledge that increased midgut permeability may also allow for extracellular ZIKV traversion of the midgut epithelium and that this is a process we did not measure in these studies. Our 0 dpf midgut permeability assays were performed within 1 hour of feeding, which could be too early to observe structural differences in the midgut epithelium. Future studies focusing on midgut integrity at higher temporal resolution are warranted.

Given that there are conserved elements of mosquito antiviral defenses and gut microbial homeostasis, including immune signaling and off-target suppression by ROS and AMPs (12,67), we were initially surprised that gene expression analyses did not detect differences in antiviral immune responses in response to antibiotic treatment. Rather, our RNA-Seq analyses confirm prior observations that show protein digestion and vitellogenesis dominate mosquito physiological processes after bloodfeeding (50), in parallel with other work showing upregulation of proteases and trypsins 2 days post-ZIKV exposure (73). Blood digestion and vitellogenesis may be hampered without microbial symbionts. Midgut symbionts may also indirectly reduce mosquito vector competence in multiple ways. These include supplementation of larval nutrition and development to increase mosquito fitness (10,74–77), stimulation of bloodfeeding behavior resulting in larger bloodmeals (although bloodmeal size differences were not observed in our studies) and increased nutrition (78), and regulation of midgut epithelial cell proliferation, where the physical microbe-midgut interface can block virus access to midgut infection (79) [although this may be pathogen specific as increased midgut infection has also been observed (20,80)]. Both microbes and ZIKV in bloodmeals resulted in increased blood digestion relative to microbe-reduced antibiotic treated mosquitoes that ingested blood without virus. It is unclear why ZIKV enhances digestion; this may result from the virus modulating cell metabolism to benefit viral replication, a pattern shown for many viruses (81). In human cells, infection with DENV elevates glycolysis to meet cellular energy needs (82). ZIKV infection of mosquito midgut cells may increase energy demand, which can be met by increasing blood metabolism. Absent ZIKV in the bloodmeal, antibiotic-treated mosquitoes exhibited digestive deficiencies during late digestion, defined here as periods beyond 2 dpf. Since blood ingestion is followed by rapid proliferation of midgut microbes that results from a heme-mediated reduction in ROS (83), the delayed digestion kinetics in antibiotic treated mosquitoes could be influenced by the disrupted microbial environment that promotes digestion.

Determining how microbes modulate the susceptibility of mosquitoes to arboviruses can help identify approaches to reduce transmission and decrease human disease. Microbial effects may also explain variability in vector competence across laboratories and geographic origins where each has different microbial communities (84), together with conventionally acknowledged factors like inbreeding during colonization (85,86). Our previous work, together with a growing body of mosquito microbiome studies, suggests that mosquito-microbe-virus interactions are environmentally variable and warrant investigation in more field-relevant contexts (15,77,87,88). Our study has several limitations. These include use of a single microbial community, ZIKV strain and dose, where disparate microbial species, community structures, and other ZIKV strains could impact susceptibility differently. Changes in microbiome composition due to antibiotic modification may differentially impact microbes with more than one copy of 16S rRNA, which could bias our evaluation of antibiotic efficacy. Unculturable bacterial species could potentially play a role in mediating hemolytic activity that could have impacted blood digestion assays. Our use of antibiotics, which incompletely eliminates microbiota, most closely represents a dysbiotic phenotype. Additionally, our gnotobiotic studies were limited to use of a single bacterial species. Newer tools including auxotrophic symbionts (89) and axenic mosquitoes (10) can be used to further investigate the metabolic roles of microbes in influencing arboviral infection of mosquitoes. The addition of ZIKV to comprise 5% of the bloodmeal volume could have resulted in augmented bloodmeal digestion compared to blood without ZIKV; mechanistic studies to determine how ZIKV impacts bloodmeal digestion are warranted. Future studies of infection dynamics at the microbe-mosquito midgut interface could prove useful in reconciling disparate vector competence outcomes and discovering novel vector control approaches.

## Acknowledgments

We acknowledge Elise Ladouceur for helping interpret H&E imaging.

**Funding Acknowledgements**:This project was funded by the Training Grant Program of the Pacific Southwest Regional Center of Excellence for Vector-Borne Diseases funded by the U.S. Centers for Disease Control and Prevention (cooperative agreement 1U01CK000516). WL was supported by the University of California, Davis, School of Veterinary Medicine Graduate Student Support Program. ALR was supported by the Coachella Valley Mosquito and Vector Control District. LLC was supported by the National Institute of Allergy and Infectious Diseases of the National Institutes of Health under award number U01AI151814. The content is solely the responsibility of the authors and does not represent the official views of the National Institutes of Health. None of the funders played a role in study design, data collection and analysis, decision to publish, or preparation of the manuscript.

## Author Contributions

Conceptualization:WL, LLC

Data Curation:WL, LKM, ETK

Formal Analysis:WL, LKM, ETK

Funding Acquisition:WL, LLC

Investigation:WL, LC

Methodology:WL, ALR, RL, LLC, GMA

Project Administration:LLC

Resources:LLC

Supervision:LLC, GMA

Validation:WL

Visualization:WL, ALR, LLC

Writing-Original Draft:WL

Writing-Reviewing and Editing:WL, ALR, LLC, LKM, ETK, GMA

---

**Figure S1.**
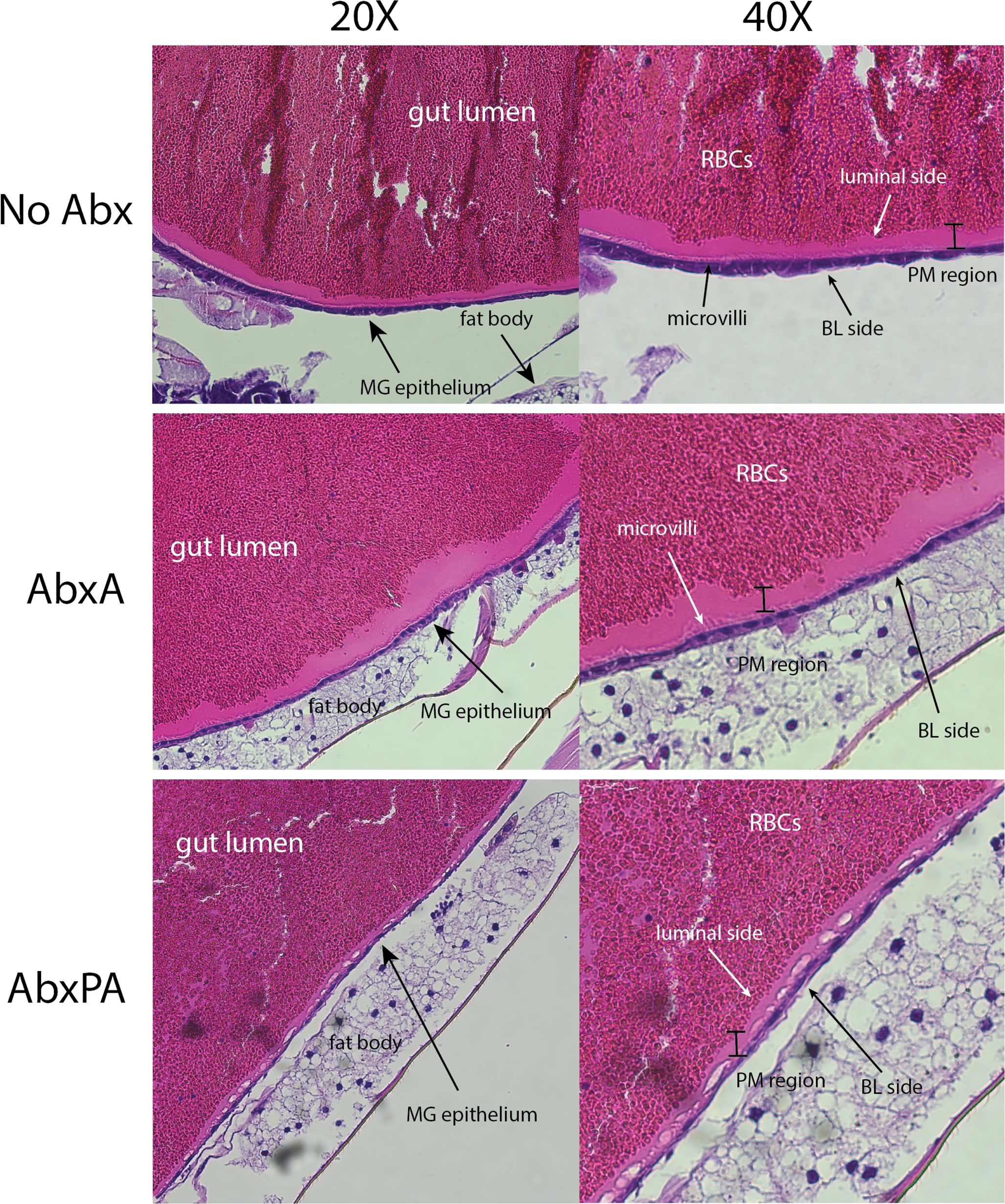
Histology of bloodfed female midguts. Cross sections of whole, bloodfed female *Ae. aegypti* along the median sagittal plane. H&E staining and bright field microscopy with a focus on the blood bolus perimeter was used to assess and identify potential structural artifacts during sample preparation that could confound immunofluorescence observations. Both 20X and 40X magnification of the same section are shown side-by-side. MG epithelium = midgut epithelium. PM region = peritrophic matrix region (where the fully formed PM would be seen). RBCs = red blood cells, from bloodmeal. BL side = basal laminal side.

**Figure S2.**
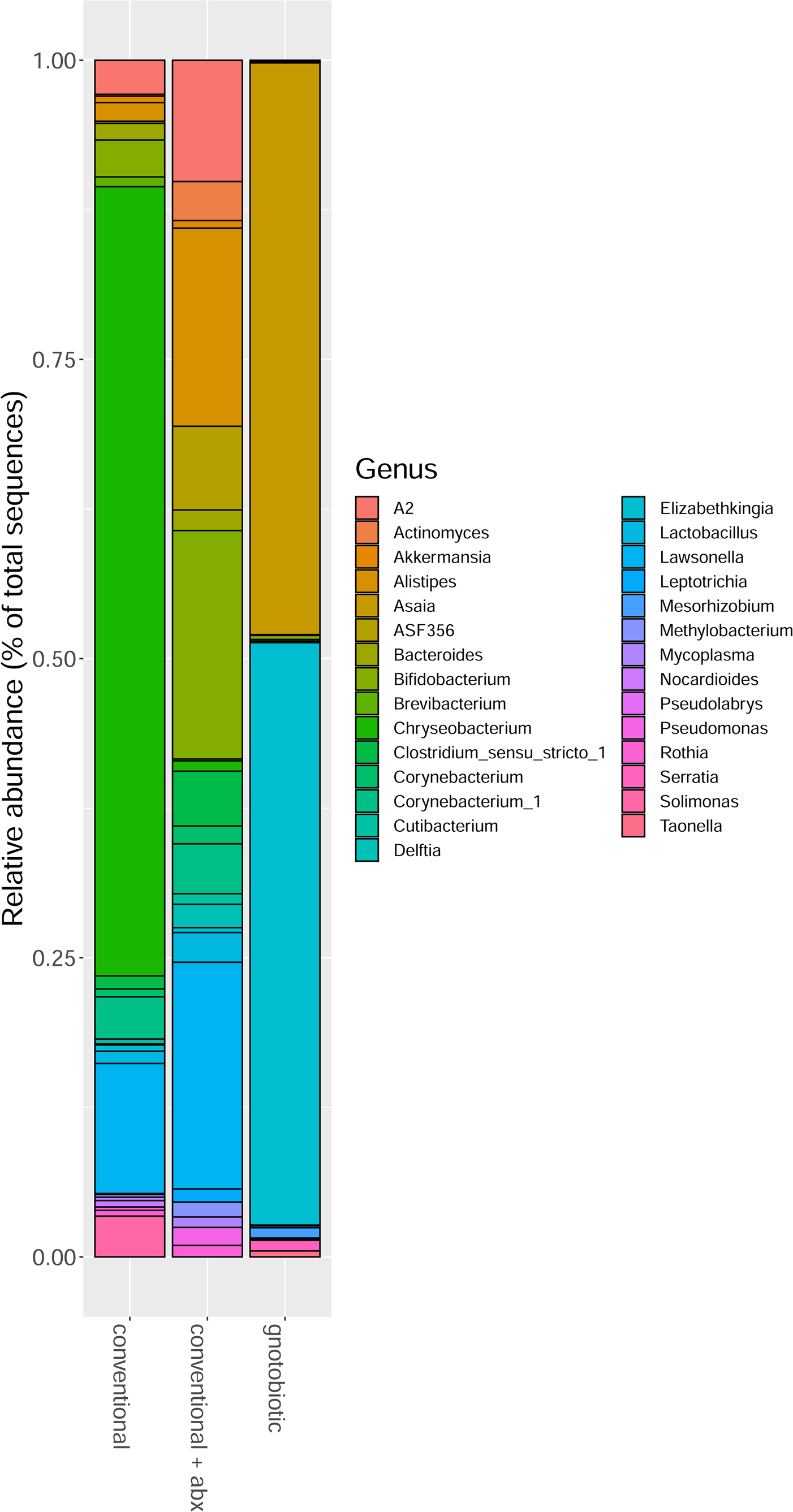
Bacterial relative abundance. Only bacteria with assigned taxonomy are shown. Each group shown represents all adult mosquitoes sequenced (N=10 Conventional, N=10 Conventional + Abx, N=8 Gnotobiotic). ASVs were agglomerated into their respective genera, represented by the colored bars.

**Figure S3.**
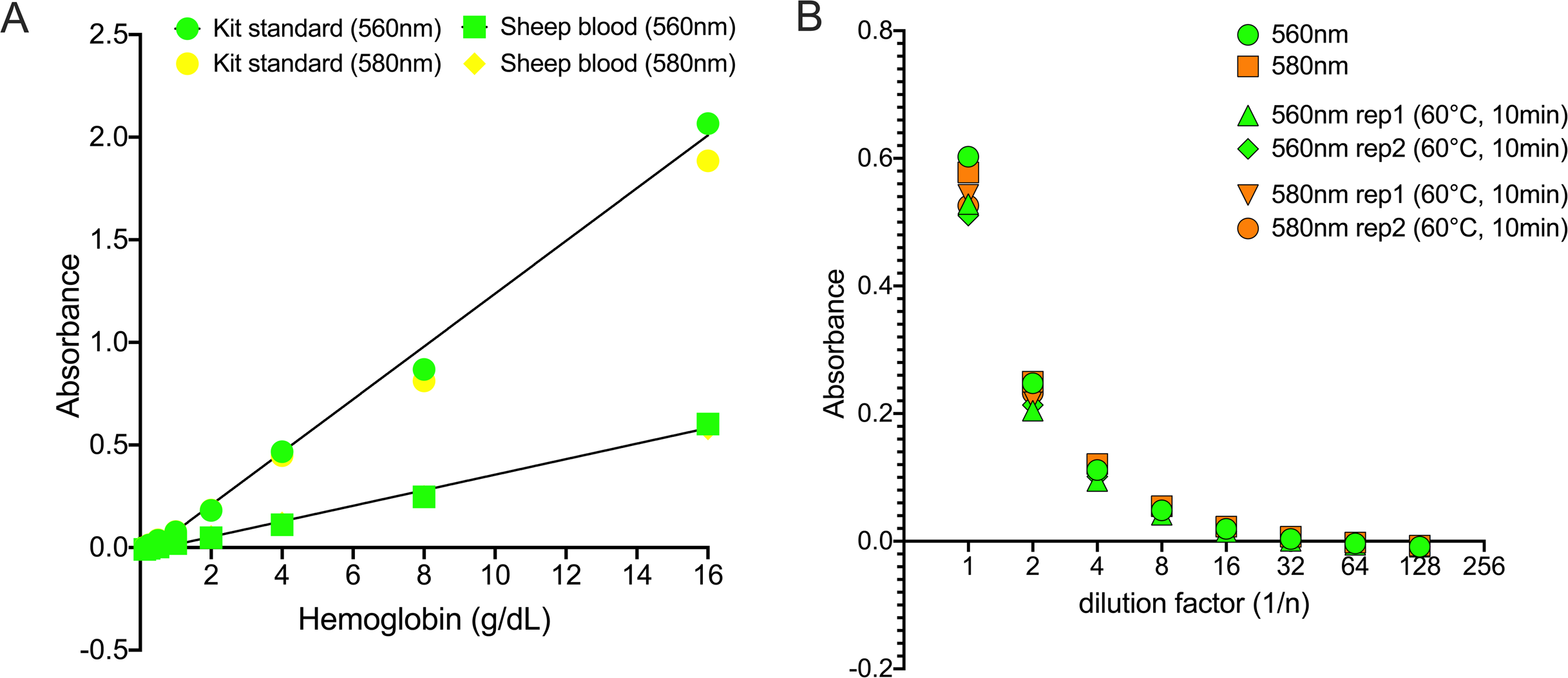
Validation of hemoglobin colorimetric assay kit for bloodfed mosquitoes. A) Serial 1:2-fold dilutions of heparinized sheep blood were tested in parallel with the kit hemoglobin standard at the absorbance extremes of 560nm and 580nm. Lines indicate standard curves generated from linear regression that were used in the hemoglobin assays. **B)** Absorbance values of sheep blood hemoglobin after heat-treating at 60°C for 10 minutes in duplicates (rep = replicate).

**Figure S4.**
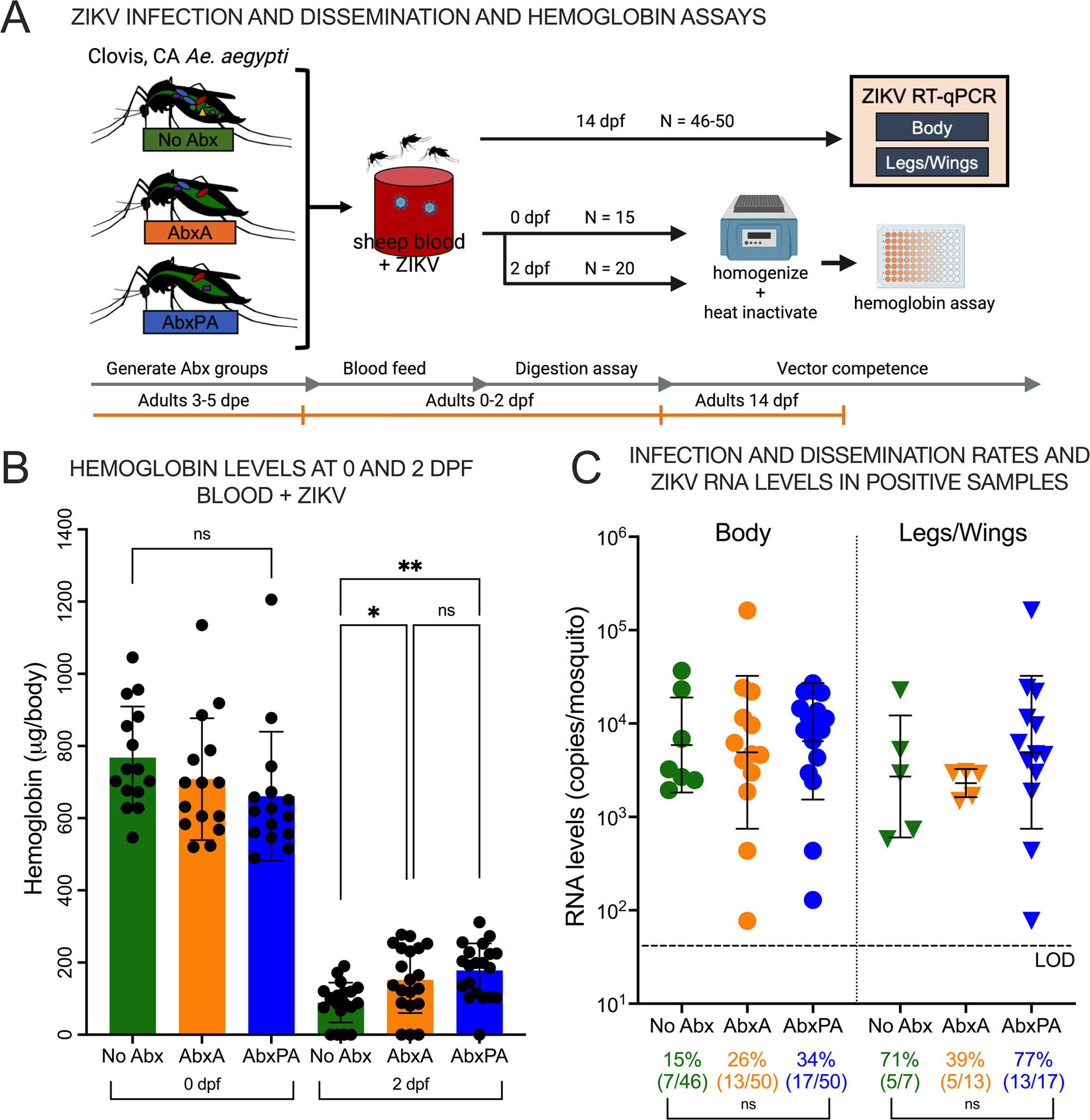
Bloodmeal digestion in microbe-disrupted *Ae. aegypti* as pupae and adults from Clovis, California. **A)** Experimental design using the same groups as in Figure 1 and hemoglobin assessments as in Figure 4 where an *Ae. aegypti* colony from Clovis, CA was used instead of *Ae. aegypti* from LA, CA. Mosquitoes were harvested 14 dpf for vector competence assessments. **B) Hemoglobin levels in *Ae. aegypti* as a proxy for bloodmeal digestion in groups that were ZIKV exposed.** Despite ingesting equivalent bloodmeal sizes, antibiotic treatment reduced blood digestion with ZIKV at 2 dpf in Clovis mosquitoes. Error bars show the means and standard deviations. N = 15 for each treatment on 0 dpf, 20 per treatment on 2 dpf. **C) Microbial disruption of pupae and adults does not impact ZIKV infection or dissemination in Clovis mosquitoes.** Rates of infection and mean ZIKV RNA levels from bloodfed Clovis *Ae. aegypti* subjected to different antibiotic treatments were not significantly different at 14 dpf. Error bars denote the geometric means and standard deviations. The dotted line denotes the average LOD which was 62 RNA copies/mosquito. ZIKV levels in bodies and legs/wings with detectable ZIKV RNA are shown in panel C. (* is P < 0.05; ** is P < 0.01; ns is not significantly different). Each symbol in B and C shows the mean value for 1 mosquito based on triplicate ZIKV RT-qPCR measurements.

**Figure S5.**
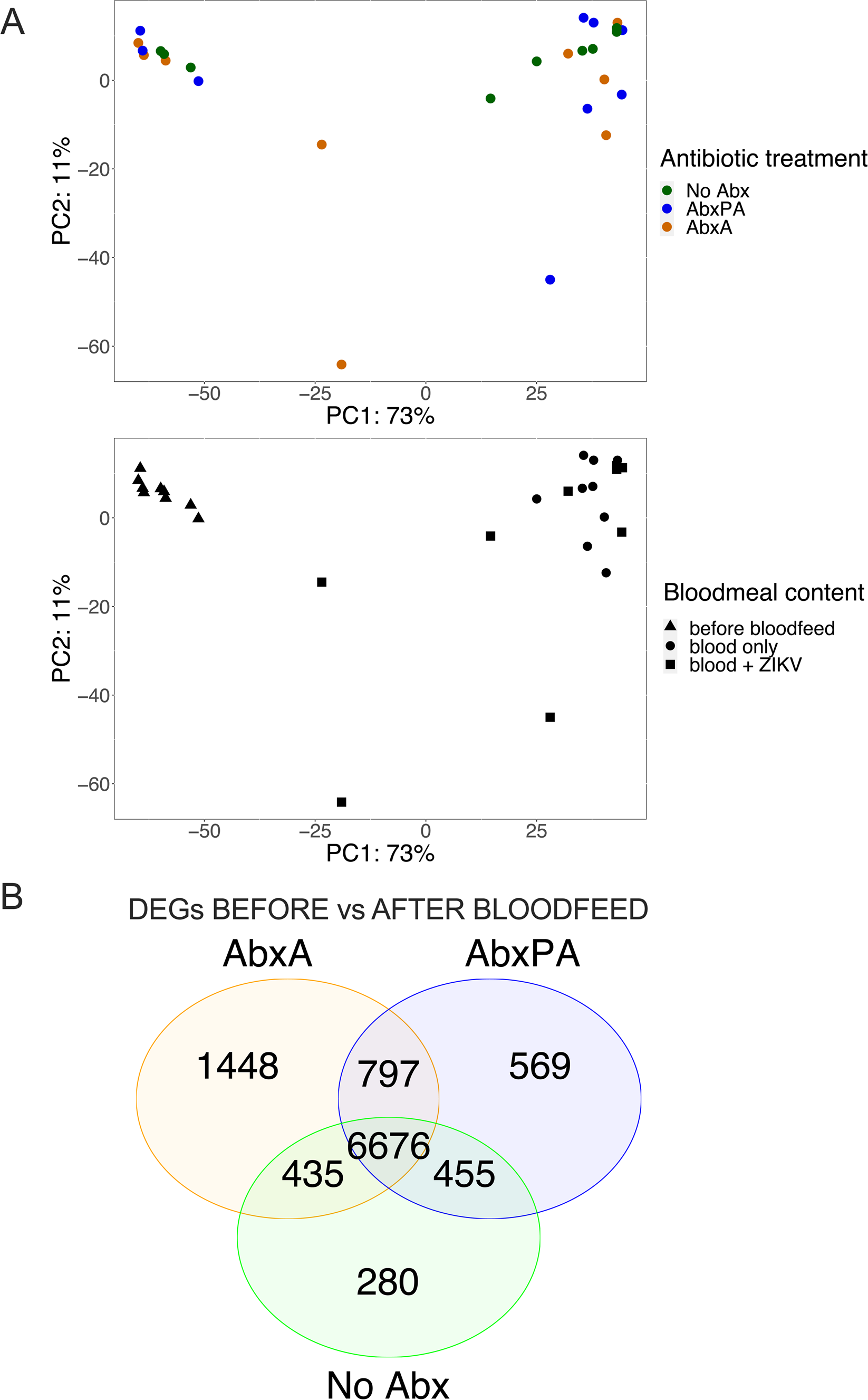
DEGs from bloodfed antibiotic-treated mosquitoes that were not exposed to ZIKV. **A)** Individual PCA plots by a single variable representing the same data in Figure 6B. Shapes indicate the bloodmeal status while colors represent the antibiotic treatments. B) **DEGs between blood-and sugar-fed *Ae. aegypti***.

**Figure S6.**
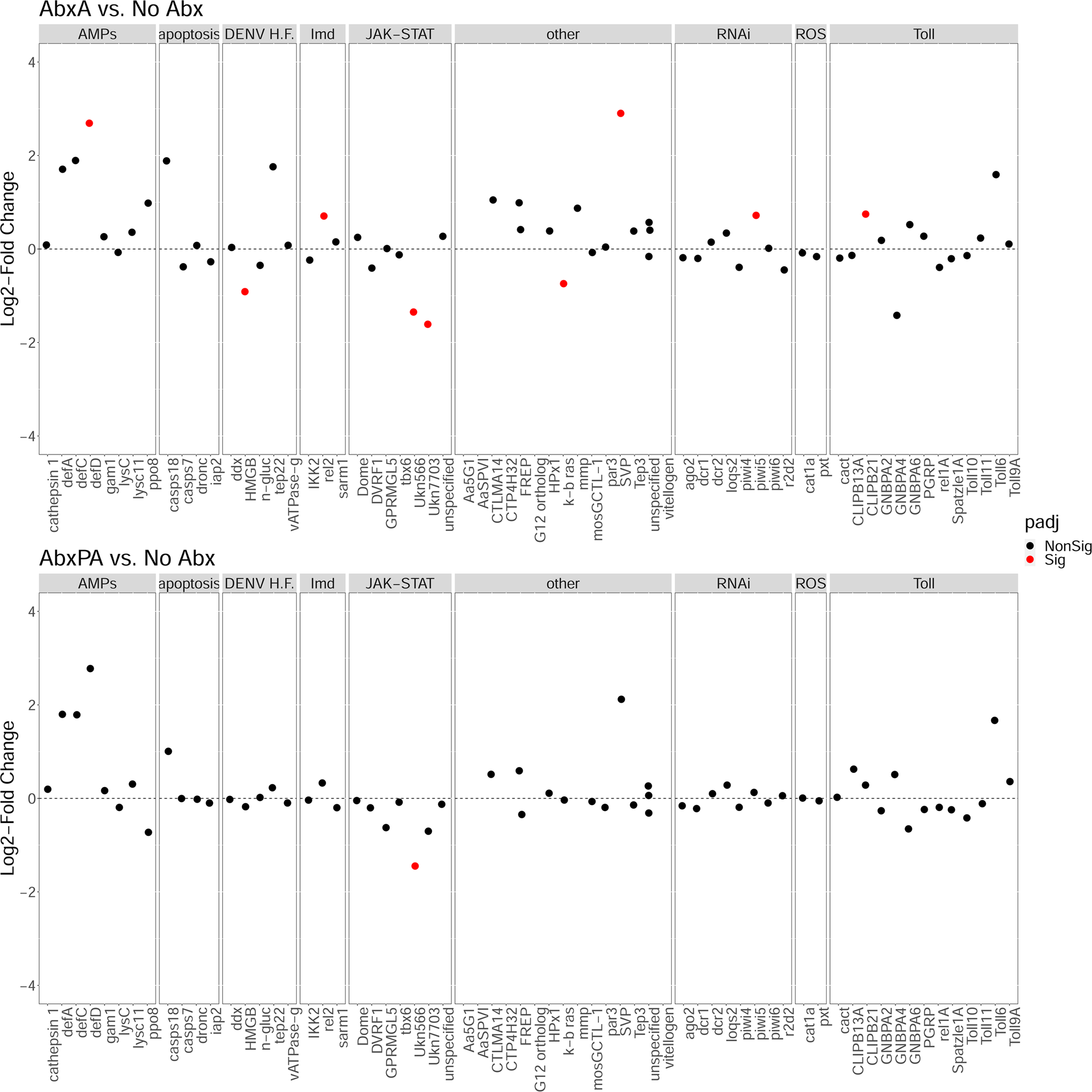
Change in immune response gene expression across antibiotic treated groups of *Ae. aegypti*. A positive log_2_-fold change indicates enrichment in AbxA or AbxPA, while a negative log_2_-fold change indicates enrichment in No Abx.

**Table S1.**
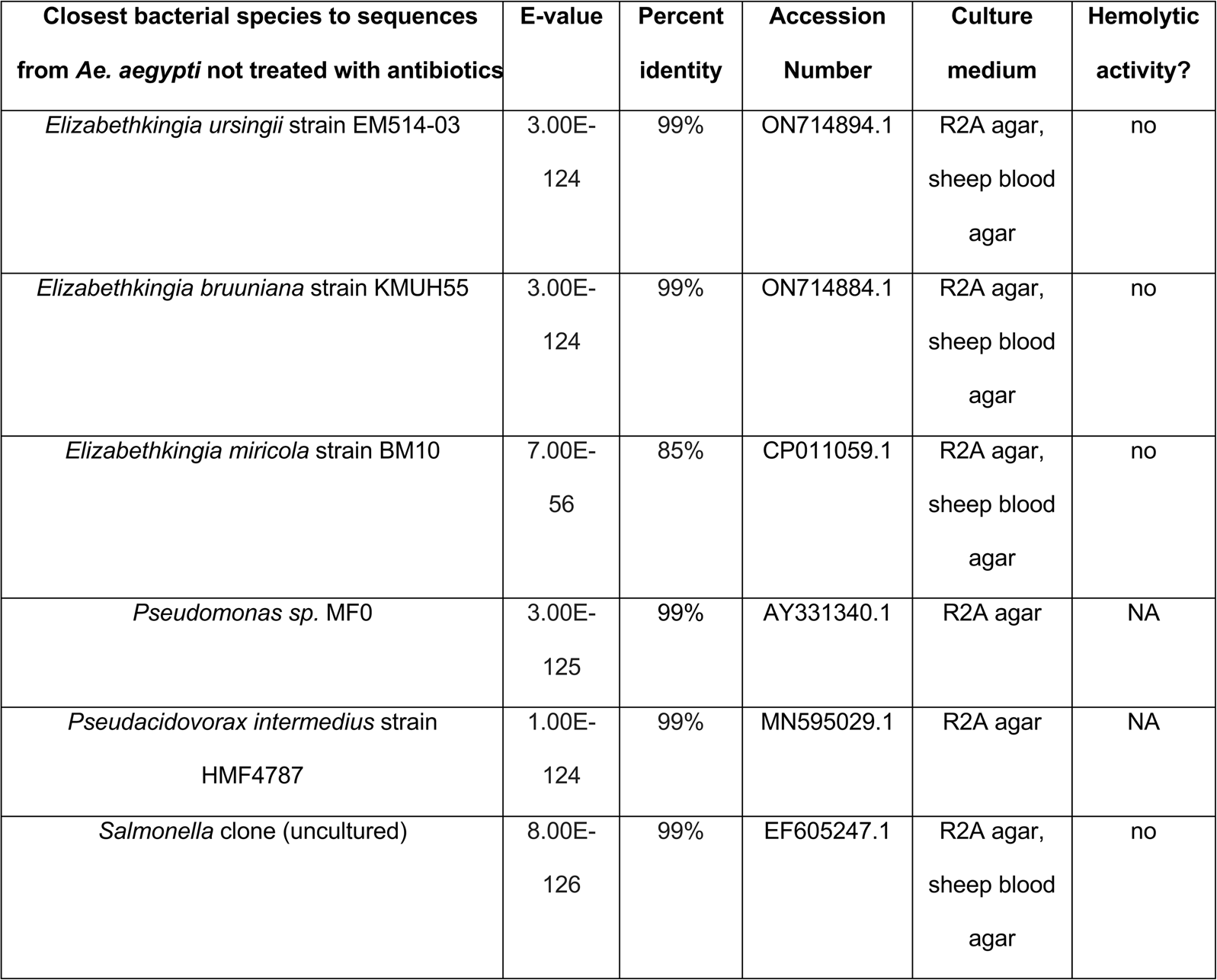
**Bacterial isolates from adult *Ae. aegypti* that were not treated with antibiotics (No Abx).** Cultured and isolated bacterial colonies from adult female *Ae. aegypti*, 3-5 dpe were Sanger sequenced at the V4 region of the 16S gene. Sequences were input into nucleotide BLAST, and the closest hits are reported. Hemolytic activity was tested by presence of ring of clearance around colonies (from lysed red blood cells) when cultured on sheep blood agar at 37°C. NA indicates hemolytic activity was not evaluated.

**Table S2.**
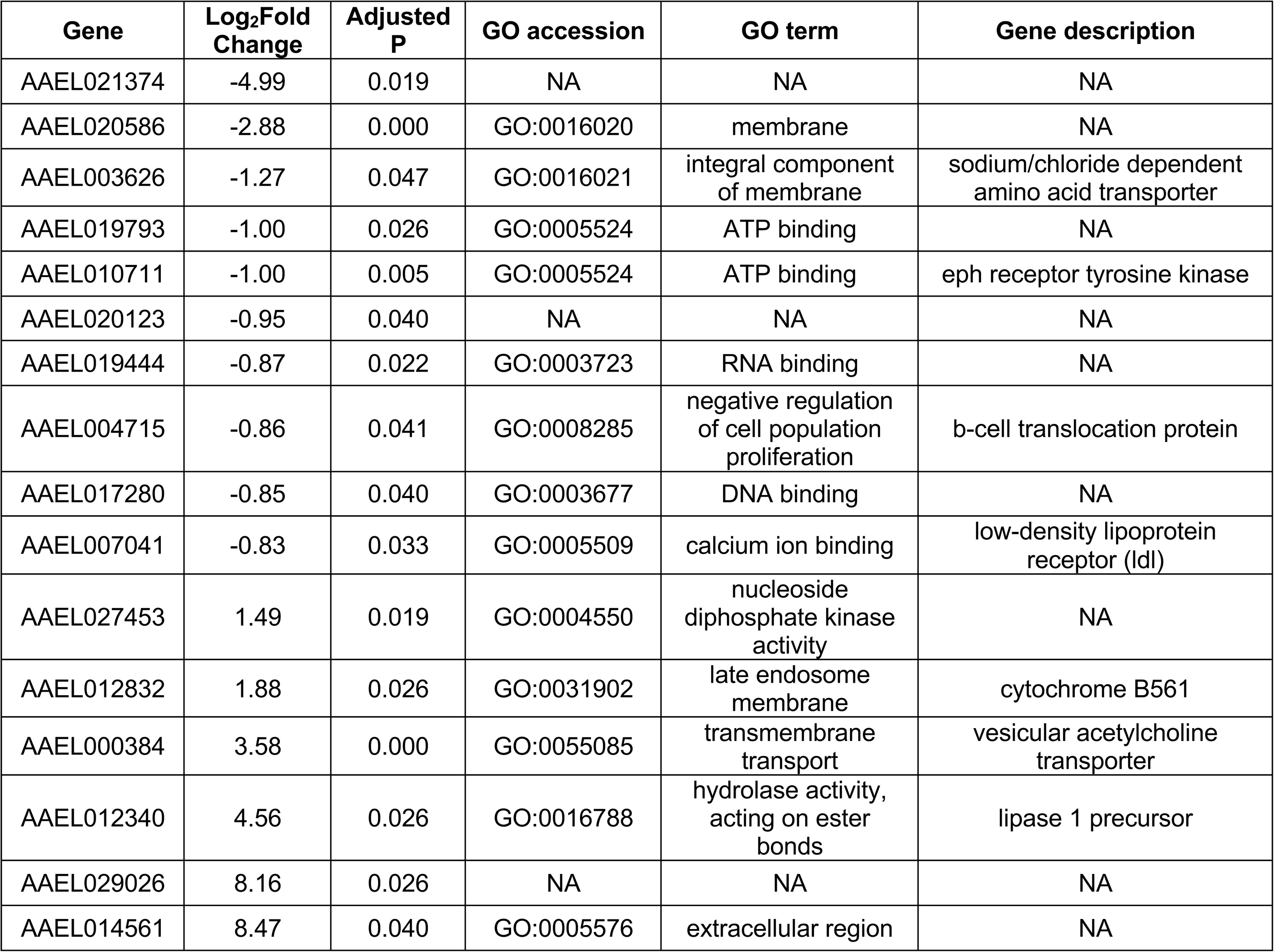
**DEGs between sugar-fed *Ae. aegypti* without antibiotics or antibiotic treated as pupae and adults.** A negative Log_2_fold-change refers to downregulation in AbxPA vs No Abx, while a positive Log_2_fold-change refers to upregulation in AbxPA vs No Abx.

| Gene | Log<sub>2</sub>Fold Change | Adjusted P | GO accession | GO term | Gene description |
| --- | --- | --- | --- | --- | --- |
| AAEL021374 | -4.99 | 0.019 | NA | NA | NA |
| AAEL020586 | -2.88 | 0.000 | GO:0016020 | membrane | NA |
| AAEL003626 | -1.27 | 0.047 | GO:0016021 | integral component of membrane | sodium/chloride dependent amino acid transporter |
| AAEL019793 | -1.00 | 0.026 | GO:0005524 | ATP binding | NA |
| AAEL010711 | -1.00 | 0.005 | GO:0005524 | ATP binding | eph receptor tyrosine kinase |
| AAEL020123 | -0.95 | 0.040 | NA | NA | NA |
| AAEL019444 | -0.87 | 0.022 | GO:0003723 | RNA binding | NA |
| AAEL004715 | -0.86 | 0.041 | GO:0008285 | negative regulation of cell population proliferation | b-cell translocation protein |
| AAEL017280 | -0.85 | 0.040 | GO:0003677 | DNA binding | NA |
| AAEL007041 | -0.83 | 0.033 | GO:0005509 | calcium ion binding | low-density lipoprotein receptor (ldl) |
| AAEL027453 | 1.49 | 0.019 | GO:0004550 | nucleoside diphosphate kinase activity | NA |
| AAEL012832 | 1.88 | 0.026 | GO:0031902 | late endosome membrane | cytochrome B561 |
| AAEL000384 | 3.58 | 0.000 | GO:0055085 | transmembrane transport | vesicular acetylcholine transporter |
| AAEL012340 | 4.56 | 0.026 | GO:0016788 | hydrolase activity, acting on ester bonds | lipase 1 precursor |
| AAEL029026 | 8.16 | 0.026 | NA | NA | NA |
| AAEL014561 | 8.47 | 0.040 | GO:0005576 | extracellular region | NA |

## Notes

https://doi.org/10.5281/zenodo.7259822

### Competing Interest Statement

The authors have declared no competing interest.

### Summary of Updates

Additional experiments to confirm gnotobiotic status performed.

